# Tracing of streptococcal strains from infant stool across human body sites links gut specificity to adhesins

**DOI:** 10.1101/2025.01.15.633122

**Authors:** Ida Ormaasen, Morten Kjos, Melanie Rae Simpson, Torbjørn Øien, Lars Snipen, Knut Rudi

## Abstract

Streptococcal species are human commensals known to colonize multiple body sites. Despite being early gut colonizers, we lack strain-level information about their origin and persistence in the gut. To gain more insight into the habitats of the streptococci present in the infant gut, we did a systematic study where mother-infant pairs were sampled from multiple body sites (stool, oral cavity, vagina, breast milk). We performed whole metagenome sequencing and isolated streptococci from 100 infant stool samples (collected at 10 days of age). To trace the streptococci at the strain level, we designed selective qPCR primers for seven streptococcal strains, and these were later utilized to screen corresponding samples from multiple body sites of the infants and their mothers. We found that two of the strains (one *Streptococcus parasanguinis* and one *Streptococcus vestibularis*) were highly prevalent in stool samples, both from infants during their first 2 years of life and from their mothers, indicating that these strains are adapted to the gut environment. Interestingly, another *S. parasanguinis* strain, closely related to the gut-prevalent strain, showed a completely different prevalence pattern, and was mainly detected in vaginal swabs, breast milk and oral swabs. Comparisons of their genomes revealed major differences in genes encoding adhesins, suggesting that host surface attachment could be a key factor for the observed differences in body site specificity. Together, our extensive tracing of streptococci across body sites of 100 infants and their mothers, provides strain-level information of prevalence patterns and reveals the presence of gut-specific streptococci.

**Importance:** Streptococci thrive on the mucosal surfaces covering the human body and colonize multiple body sites. To determine the distinct streptococcal composition in each habitat and to evaluate their presence in different habitats, strain-level characterization is crucial. We show that two closely related strains, both isolated from stool, are distributed differently across the human body, with one of them prevalent in the stool samples and the other more prevalent in other samples. This emphasizes the necessity of strain-level analysis for the identification of true colonizers of a habitat.

## Introduction

After birth, the infant’s body undergoes an immediate bacterial colonization. While most colonizers are transient, other persist (1, 2). The *Streptococcus* genus is a diverse group of both beneficial and harmful species that can colonize multiple sites of the human body (3). Shortly after birth, streptococci colonize the infant gut (4-6), with high abundance the first weeks (7-9). These bacteria remain as members of the gut microbiota throughout life, although their relative abundance significantly decreases after a few months post-delivery (10-12). Studies suggest that streptococci can be transmitted to the infant gut from maternal reservoirs (13-15). Although mechanisms for gut-bacterial sharing early in life have been studied (16), we still lack knowledge about the gut microbiota establishment at the strain level. To identify gut-colonizing streptococci and their potential maternal reservoirs, it is crucial to trace individual isolates and further characterize these for phenotypic traits that can explain the colonization.

More than 30 streptococcal species have been isolated from the human body (17). Although some are pathogenic (18), most species are commensals (17, 19) or even display probiotic effects (20, 21). As facultative anaerobes, streptococci obtain energy by fermenting simple sugars such as glucose and lactose into lactate, both in the presence or absence of oxygen (22). Such versatility is important, since oxygen is depleted in the infant gut the first days after birth, leading to an anaerobic gut environment and subsequent establishment of obligate anaerobes (4). Some studies indicate that streptococci may have a key role in the adult gut, more specifically in the small intestine. It has been reported that *Streptococcus* is one of the most abundant genera in the ileal microbiota (11, 12, 23), and omics analyses of the ileal streptococci and their metabolic activity imply that these species are involved in the carbohydrate degradation that takes place in the small intestine (19, 24, 25). Considering that streptococci may have an important role in the small intestine microbiota, more knowledge on the gut streptococci is needed.

Streptococci are commonly found in the oral cavity, upper respiratory tract, skin and genitourinary system in addition to the gut (26). The oral cavity is colonized by an abundant streptococcal population (27). Reported members of the oral microbiota include *Streptococcus parasanguinis, Streptococcus gordonii, Streptococcus salivarius* and *Streptococcus vestibularis*. The three latter species are also associated with the gastrointestinal tract (28-30). Other common human streptococcal species include *Streptococcus pneumoniae* in the upper respiratory tract (31) and *Streptococcus agalactiae* in the genitourinary system (32). Moreover, various streptococcal species like *S. parasanguinis, S. salivarius and S. vestibularis* have been found in breast milk (33-35). Different genetic and environmental factors probably influence the body site preference of streptococci, including nutrient availability. Importantly, streptococci produce various adhesins used in host surface attachment (26). These cell surface-associated proteins recognize and bind to receptors found in the different tissues on the human body (36). Therefore, the colonization sites depend on the adhesin targets, in addition to environmental conditions like oxygen and pH levels (26).

Given the widespread presence of the *Streptococcus* genus across the human body, strain-level characterization of the streptococcal populations in the different habitats is crucial. Whole metagenome sequencing has made it possible, to some degree, to characterize the most abundant bacterial species in a metagenome at the strain level, making it suitable for determining prevalence and abundance of specific strains in a collection of samples (37). However, further phenotypic characterization of strain-specific traits, beyond what metagenomics can provide, demands possession of strain isolates (38). Furthermore, the possession of strain isolates enables reconstruction of their genomes, which again allows large scale tracing of strains by sensitive and low-cost methods such as real-time quantitative PCR (qPCR) screenings (39, 40). This underlines the importance of bacterial cultivation in combination with metagenomic analyses.

Here, we investigated the prevalence of streptococcal isolates across different body sites in 100 mother-infant pairs from the Probiotics in Prevention of Allergy among Children in Trondheim (ProPACT) study (41). The experimental setup is illustrated in Fig. 1. By performing whole genome sequencing of streptococcal isolates derived from infant stool samples collected at 10 days of age, combined with whole metagenomic sequencing of the same samples, we designed primers specific for each strain among the isolates. The strain-specific primers allowed us to perform a comprehensive qPCR-based screen of different samples from the mothers (stool, vagina, breast milk) and their infants (stool, oral cavity), aiming to determine the overall prevalence of the streptococcal strains. Firstly, we present the relative distribution of seven streptococcal strains, in which two are prominent gut-associated strains. Secondly, we suggest that the distinct habitat pattern observed for two closely related strains is partly due to difference in proteins related to host surface attachment.

**Fig 1.**
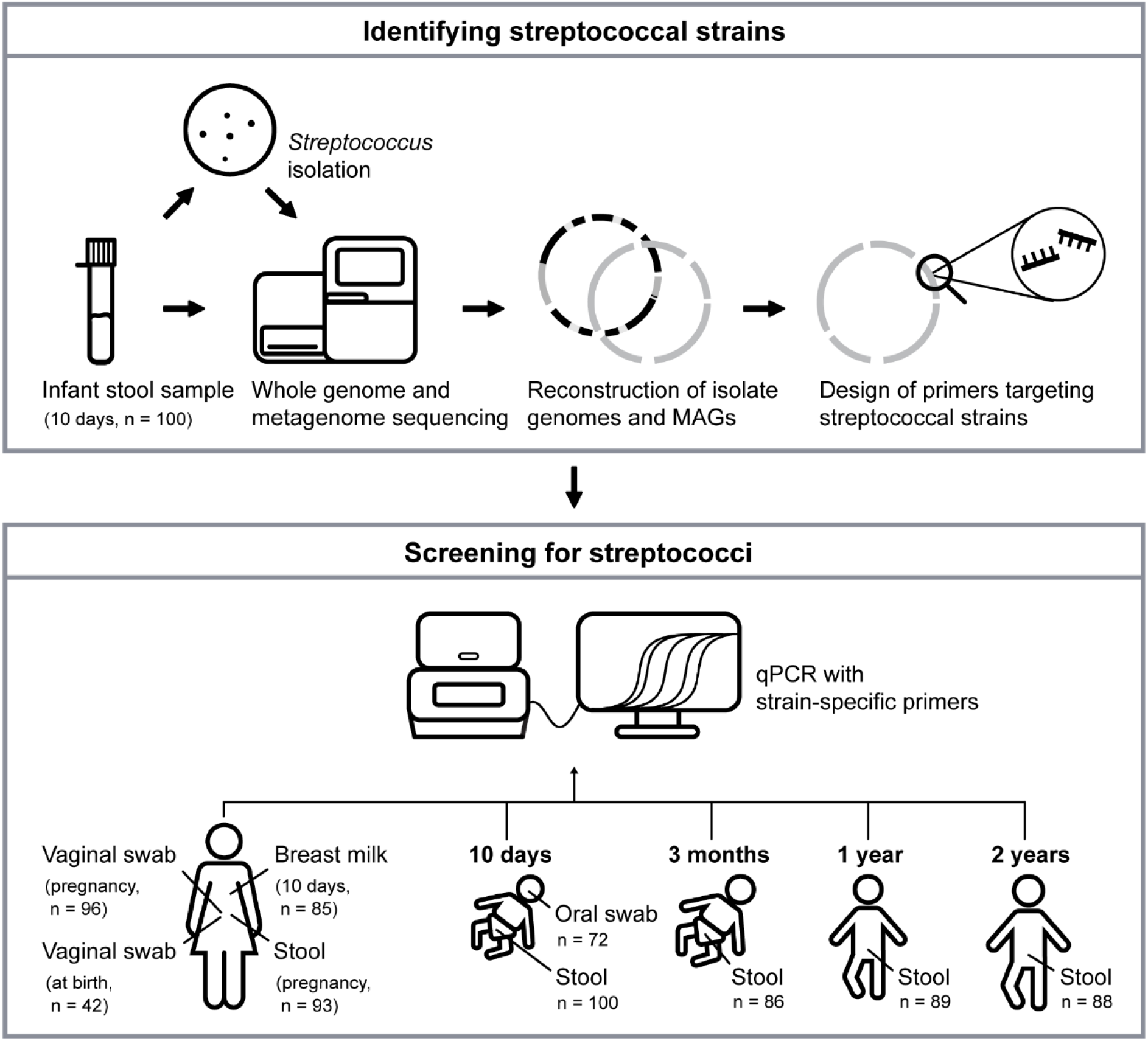
Experimental setup of the streptococcal isolation and screening. Stool samples, collected from 10-day-old infants (n=100), were cultivated on streptococcal-selective agar plates to obtain streptococcal isolates. Whole metagenome sequencing of the stool samples and whole genome sequencing of the isolates resulted in the reconstruction of the isolate genomes and streptococcal metagenome-assembled genomes (MAGs). These were used in the design of primers that are specific to each of the streptococcal strains isolated from the stool samples. The designed primers were applied in a qPCR screening to enable the detection of the streptococcal strains and subsequent tracing of these across samples from the infants and their mothers (stool, oral swabs, vaginal swabs, breast milk). The different samples had been collected in week 30 of pregnancy and at birth, as well as 10 days, 3 months, 1 year and 2 years after birth, as shown in the illustration.

## Results

### The overall distribution of *Streptococcus* in the different sample types

To investigate the presence of streptococci in the collection of samples, we analyzed previously generated 16S rRNA gene sequencing data (41-44). The analysis identified *Streptococcus* species in all the sample types (Fig. 2). Overall, the highest average relative abundances of streptococci were found in the infant oral cavity (79%) and breast milk (50%). In the infant stool samples, the highest average relative abundance was observed in the stool collected at 10 days of age (12%), followed 3 months samples (6%). In the stool samples collected at 1 year and 2 years of age, the average relative abundances had decreased to 3% and 2%, respectively.

**Fig 2.**
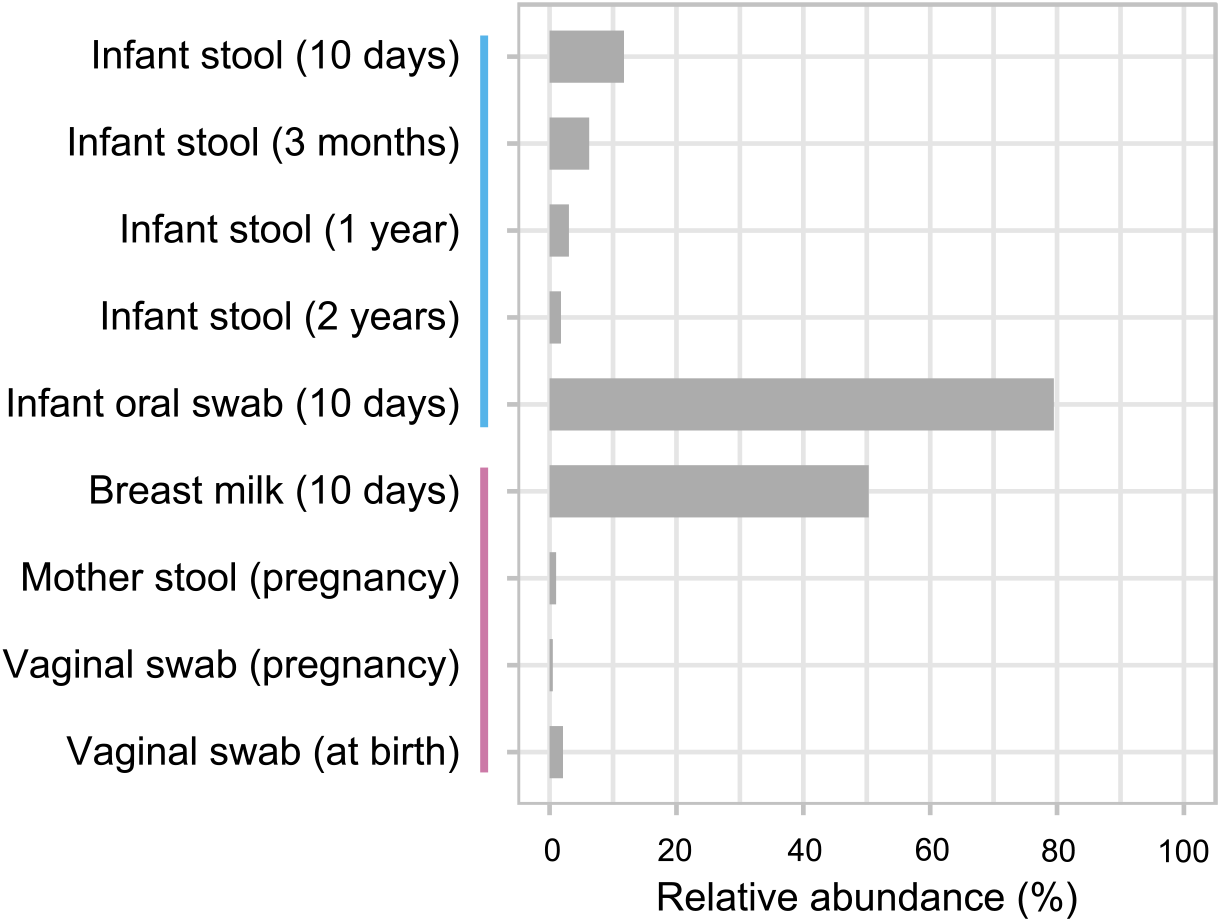
Relative abundance of *Streptococcus*. The figure shows the average relative abundance of Streptococcus in the different sample types, determined by 16S rRNA gene analysis. The time points of sample collection are given in parentheses. The vertical lines on the y-axis indicate the samples from the infants (blue) and the mothers (pink).

### Streptococcal diversity in the infant gut microbiota at 10 days of age

Since the 16S rRNA gene analysis revealed that the highest average streptococcal relative abundance in the infant stool samples was at 10 days of age, we selected these for streptococcal-selective cultivation. Out of the 100 samples, 69 samples displayed growth on the streptococcal-selective agar. A total of 177 bacterial isolates were obtained from 53 different samples. By Sanger sequencing of the full length 16S rRNA gene, the most abundant genera among the isolates were identified as enterococci (56%), lactobacilli (18%) and streptococci (9%). Further taxonomic classification of the streptococcal isolates (n = 16) was achieved by whole genome sequencing. This revealed that the isolates belonged to seven species, as defined by GTDB taxonomy: *Streptococcus parasanguinis_D* (n = 2), *Streptococcus parasanguinis_I* (n = 2), *Streptococcus parasanguinis_F* (n = 1), *S*. sp900766505 (n = 3), *Streptococcus agalactiae* (n = 5), *Streptococcus vaginalis* (n = 2), and *Streptococcus vestibularis* (n = 1) (Table S1).

Complementary to the cultivation-based approach, we performed whole metagenome sequencing of the same 100 infant stool samples and subsequent assembly, binning and genome reconstruction to identify *Streptococcus* spp.. The sequencing resulted in 112 metagenome-assembled genomes (MAGs), of which 14 were classified as streptococci (Table S2). Among the streptococcal MAGs, several represented the same species as the isolates (*S. parasanguinis_D* (n = 1), *S. agalactiae* (n = 1), *S*. sp900766505 (n = 1), *S. vaginalis* (n = 1), and *S. vestibularis* (n = 1)). The remaining streptococcal MAGs were assigned to the following species according to the GTDB taxonomy: *S. parasanguinis_S* (n = 1), *S. salivarius* (n = 1), *S. infantis_M* (n = 1), *S. lactarius* (n = 1), *S*. sp001556435 (n = 1), *S*. sp000187445 (n = 1), and *Streptococcus* spp. (n = 3). To investigate the phylogenetic relationship between the streptococcal isolates and MAGs, we determined the estimated evolutionary distances between them. As a reference, we also included complete genomes from the GTDB database that belonged to the same species as the isolates and MAGs. The estimated evolutionary distances, illustrated in the dendrogram in Fig. S1, show that the isolate assemblies cluster according to their assigned species. The same can be seen for most of the MAGs. Furthermore, some of the isolates, including *S. vaginalis* isolate 14, *S. agalactiae* isolate 13, *S. parasanguinis* isolates 8 and 6, and *S. parasanguinis* isolates 5 and 3 (Fig. S1) seem to be identical to a MAG within the same cluster, since the corresponding estimated evolutionary distances are close to zero. Based on the estimated evolutionary distances between the 16 isolates, we concluded that these represent nine different strains, including two *S. agalactiae*, three *S. parasanguinis* (*S. parasanguinis_D, S. parasanguinis_I* and *S. parasanguinis_F* according to GTDB taxonomy), one *S*. sp900766505, two *S. vaginalis* and one *S. vestibularis*. These will be referred to as strains below. To distinguish between the strains belonging to the same species, a number was assigned to their names (Table S1).

We investigated how abundant the nine streptococcal strains and the streptococcal MAGs were across the infant stool samples at 10 days of age (see Fig. S2 for total bacterial composition in the infant stool samples). On average, the relative abundance of streptococci in these samples, determined by whole metagenome sequencing, was 6.3%, and the highest occurring abundance in a sample was 73.3% (Fig. 3). Both the strains and the streptococcal MAGs not represented by any of the strains contributed substantially to the relative abundance in the samples (Fig. 3). Notably, the *S. vestibularis* strain was substantially more prevalent and abundant than the other strains, contributing to at least 1% of the bacterial composition in 28% of the samples (Fig. 3).

**Fig 3.**
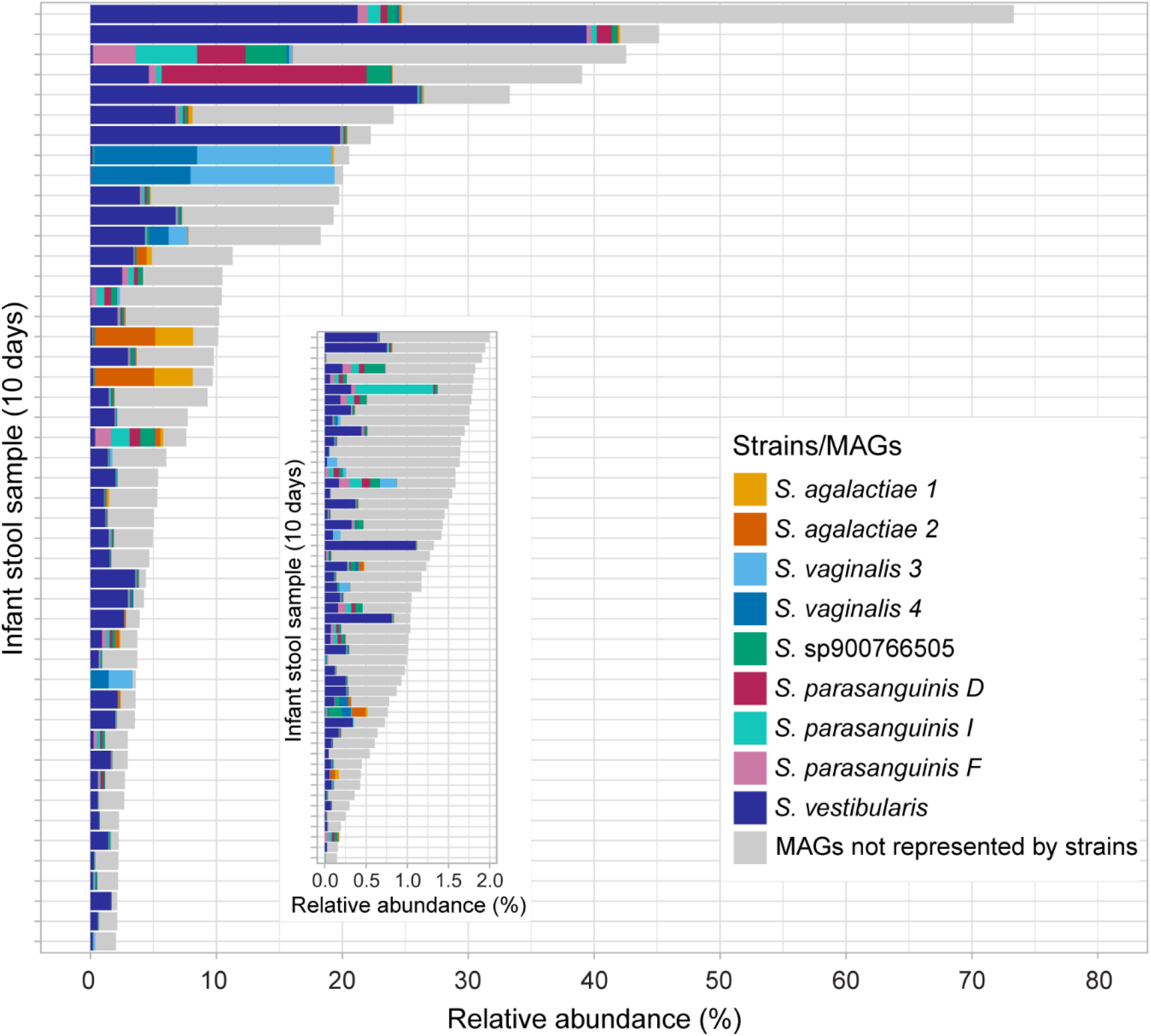
Streptococci in the infant gut at 10 days of age. The relative abundance of streptococci, as determined by whole metagenome sequencing, is shown. The relative abundances of the nine streptococcal strains in the infant stool samples are indicated in colors. The relative abundances of the streptococcal MAGs with a different taxonomic classification than the nine strains are shown in grey. Note that the stool samples with a streptococcal relative abundance less than 2% are shown in a different scale.

### Streptococcal screening of infant- and mother-derived samples reveals strain-specific prevalence patterns

To study the prevalence of the strains across the samples collected from the mother-infant pairs, we performed a screening with strain-specific primers (Fig. 1). The primers were thoroughly designed to target nucleotide sequences unique to each strain. Systematic testing of the primers showed that 7 out of the 9 designed primers could successfully be used to specifically identify the targeted strain (we were not able to make strain-specific primers for the strains *S*. sp900766505 and *S. parasanguinis D*). These seven primer pairs were used further in the qPCR-based tracing of strains. The results from the primer design and testing can be found in Table S3.

In line with the metagenome data (Fig. 3), the qPCR screening for the streptococcal strains across the samples revealed that the *S. vestibularis* strain was by far the most prevalent strain in the infant stool samples collected at 10 days of age, being detected in 84% of the samples (Fig. 4). Further, the *S. vestibularis* strain was frequently detected in the infant oral samples (44%), the breast milk samples (32%) and the mothers’ stool samples (44%) (Fig. 4), making it the most prevalent strain across the whole sample collection (27%). The second most overall prevalent strain was *S. parasanguinis F*, which was found to be present in 19% of the sample collection. This strain was detected across most of the sample types, but it was particularly prevalent in breast milk (53%) and vaginal samples (pregnancy) (51%) (Fig. 4). In contrast, the closely related strain *S. parasanguinis I* displayed a completely different detection pattern than *S. parasanguinis F* (Fig. 4). *S. parasanguinis I* was detected mainly in stool samples, both in mothers’ stool and infant stool collected at 10 days, 1 year and 2 years of age (38%, 14%, 35% and 19%, respectively). When it comes to the two *S. vaginalis* strains, they shared a similar pattern of detection across the samples (Fig. 4). They were detected mainly in mothers’ stool samples, vaginal samples during pregnancy and infant stool samples (10 days). The two *S. agalactiae* isolates were the least prevalent strains in the screening, with only six detections in total.

**Fig 4.**
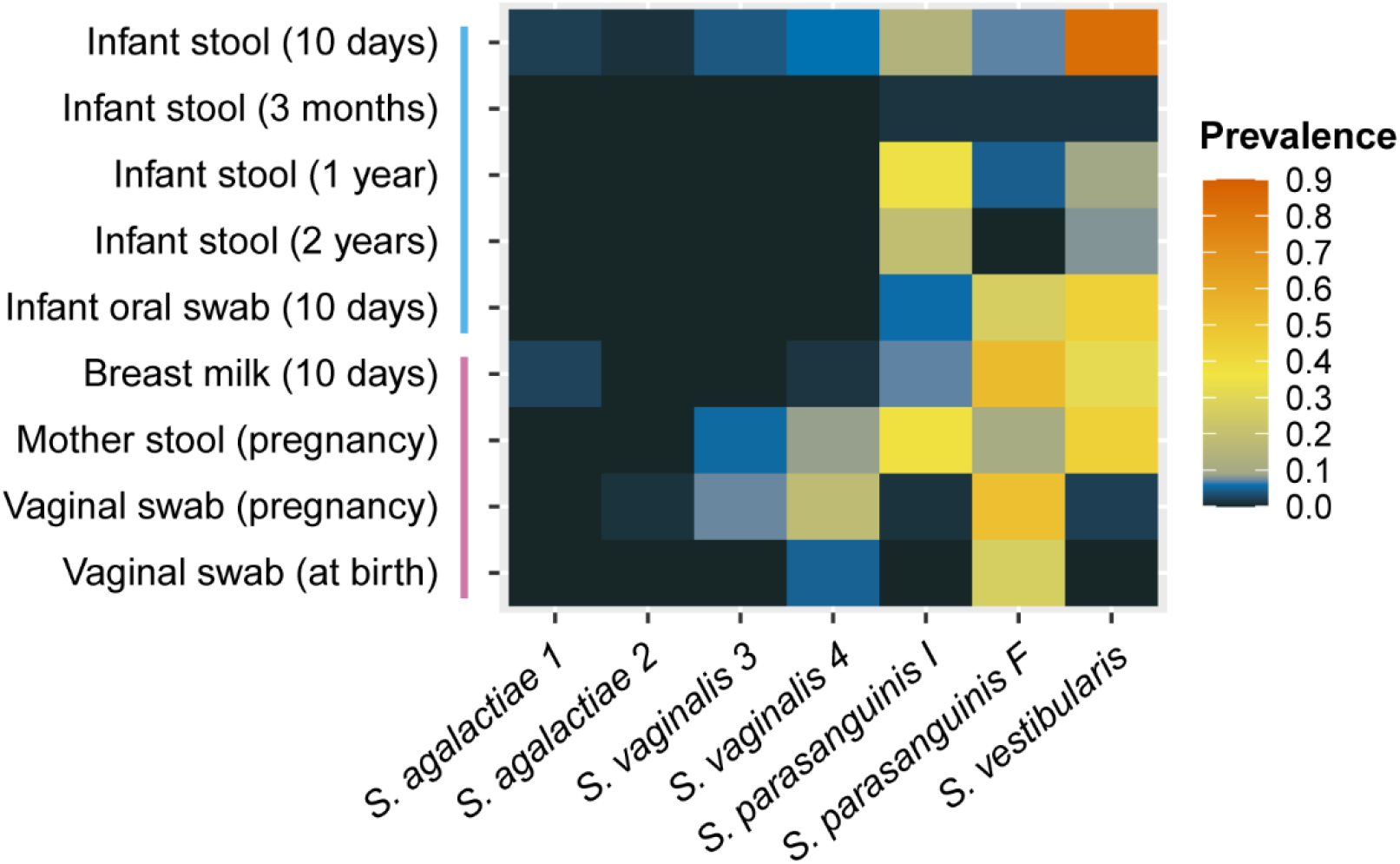
Strain prevalence. Prevalence of the strains in all the samples from maternal and infant body sites at several time points, based on a qPCR screen with strain-specific primers. The time points of sample collection are given in parentheses. The vertical lines on the y-axis indicate the samples from the infants (blue) and the mothers (pink). A detailed overview of the strain detections in each sample is given in Fig. S3.

Detections of the same strain in both infant- and mother-derived samples within one mother-infant pair were frequently made (Fig. S3). There was a statistically significant association between *S. vestibularis* detection in the infant oral cavity and breast milk (Fisher’s test, p < 0.005). The odds ratio of the statistical test indicated that it was 10.8 times more likely that *S. vestibularis* was detected in the breast milk if it was detected in the infant oral cavity as well. When testing for associations with strains detected in infant stool samples, no statistically significant associations were found. The p-values and odds ratios for all tested associations can be found in Table S4.

### Phenotypic and genotypic comparison of the strains

To investigate whether the genetically different strains also exhibited phenotypic distinct traits, their capability to utilize various substrates, including carbohydrates, was tested using the Rapid ID 32 Strep test. Although many of the strains displayed enzymatic activity towards the same substrates, there were differences in substrate utilization among the strains, even between the strains belonging to the same species or closely related species (Fig. S4). The differences seen for the closely related *S. parasanguinis F* and *I* strains were the latter’s additional capability to utilize D-ribose D-mannitol, Dtrehalose, D-melibiose, pullulan, resorufin-ßD-glucopyranoside and methyl-ßDglucopyranoside. Among all the strains, *S. vestibularis* was the only strain capable of utilizing urea.

The strikingly opposite prevalence patterns of the two closely related *S. parasanguinis* strains prompted us to further investigate these genomes in more detail. We compared their genomes to identify dissimilar genes (see Methods). We anticipated some gene hits (instances where a query from the one strain aligns, to some degree, with a position on a contig from the other strain) to be the same, regardless of the strain used as query (whether *S. parasanguinis I* was blasted against *S. parasanguinis F* or vice versa). The comparison resulted in 28 gene hits for *S. parasanguinis I* and 32 for *S. parasanguinis F*, in which 16 of the hits were the same for both strains (Table S5). Notably, 6 of these 16 hits were genes that encode surface proteins and putative adhesins potentially involved in colonization (Table S5). In addition, two adhesin genes were only found in the *S. parasanguinis I* strain, and three other adhesin genes were detected solely in the *S. parasanguinis F* strain. A closer inspection of the sequence alignments and domain organization in these putative adhesins (11 in total) revealed sequence differences and variable numbers of domain repeats (Table S6). For example, we found dissimilarities in surface attachment-related protein domain regions in the genes encoding the GbpC/Spa domain-containing protein, the SspB-related isopeptide-forming adhesin and the CshA/CshB family fibrillar adhesin-related protein, as illustrated in Fig. S5.

## Discussion

Streptococcal species are known to colonize a number of body sites; however, strain-level information is mostly lacking. Here we performed a large-scale tracing of individual streptococcal strains isolated from stool samples of 10-day-old infants across body sites of 100 mother-infant pairs. Most prominently, this strain-level mapping revealed gut-adapted streptococcal strains, suggested that the maternal gut microbiota serves as a reservoir for these, and pointed towards genes related to host surface attachment as one of the key drivers for the body site preference of streptococcal strains.

The two strains *S. parasanguinis I* and *S. vestibularis* were highly prevalent in the infant stool samples collected at different time points up to 2 years. Our strain-specific tracing revealed that these strains were also the most prevalent in the stool samples of the mothers, indicating that they are adapted to the human gut environment. In line with these results, Ferretti et al. also found that bacterial strains present in mothers’ stool samples were more persistent in the infant gut compared to strains from other sample types (15). Together with our results, this strongly suggest that the maternal gut microbiota serves as an important reservoir for gut-colonizing streptococcal strains in the infant. The ecological niches of streptococci in the gut are not well elaborated, but streptococci have been suggested to thrive within the small intestine (11, 19, 24, 25). We specifically looked for the *S. parasanguinis I* and *S. vestibularis* strains in stool samples, however, their representative species have been found in effluents from the small intestine (19, 24). This indicates that these strains might be able to occupy niches in the small intestine.

The two closely related strains *S. parasanguinis I* and *F* displayed distinct prevalence patterns across the samples from different body sites (Fig. 4). This indicates that despite their genetic similarity, these strains colonize different habitats. *The S. parasanguinis I* strain occurred mainly in the stool samples, while the *S. parasanguinis F* strain was frequently found in the other sample types, including the infant oral samples. Recognized as a common member of the oral microbiota (27), the species *S. parasanguinis* has been speculated to appear in the gut due to a translocation from the mouth (45), in which the oral-adapted species only transiently appears (46, 47). Our finding suggests that there is a more nuanced explanation for *S. parasanguinis* presence in the gut than previously proposed, involving gut colonization of specific *S. parasanguinis* strains.

The habitat differences seen for the *S. parasanguinis I* and *F* strains were further investigated by substrate utilization tests and by searching for strain-specific variation in their genomes. The two strains displayed differences in substrate utilization capabilities, and this may be a contributing factor to the habitat preferences of the strains. Moreover, the genome comparison also revealed that a major fraction of the genetic variation between the strains were found in genes encoding adhesins or adhesin-associated proteins, such as the GbpC/Spa domain-containing protein, the SspB-related isopeptide-forming adhesin and the CshA/CshB family fibrillar adhesin-related protein (Table S6). Genetic variations were found in regions encoding protein domains involved in surface attachment. An example of this was observed in the GbpC/Spa domain-containing protein, where the number of repeats in the cell surface antigen I/II A repeat domain varied. Streptococci are known to produce a great variety of adhesins (48), such as the antigen I/II (AgI/II) polypeptide family adhesins (49). Orthologous AgI/II proteins are found throughout the *Streptococcus* genus. Species-specific variations among these include different number of repeats in the alanine-rich repeats (A region) and the proline-rich repeats (P region), as well as dissimilarities in the flanking variable region (V region) (49, 50). Adhesin genes from streptococcal strains within one species have been reported to display high sequence conservation (51, 52). Based on this, it has been suggested that the divergence seen in orthologous AgI/II adhesins is due to niche adaptations evolved in the various species (49, 50). In contrast, our results showing differences in the prevalence patterns between *S. parasanguinis I* and *S. parasanguinis F* suggest that distinct sets of adhesins used for attachment to host surfaces may also play an important role for strain-level variation in colonization.

Three out of seven strains in the screening were not detected in the samples from which they were originally isolated, demonstrating that the strict criteria affected the strain detection sensitivity. This is partly caused by the chosen study design, in which strict terms were set for the screening analysis to ensure high specificity. Furthermore, the streptococcal-selective cultivation resulted in the revival of only 16 streptococcal isolates, belonging to nine strains. Although the whole metagenome sequencing identified additional species, the distribution of the streptococcal strains and MAGs in the samples shows that the isolated strains are representative for the streptococcal population in the infant gut 10 days after birth in our study population. Notably, while the same strains were also detected in the infant stool after 1 and two years, few detections of these strains were made in the infant stool samples collected at 3 months of age. The 16S rRNA gene sequencing results showed that *Streptococcus* was present in the gut at this time point. The reason for this discrepancy is not known. A possible explanation for this may be that the strains found to be prevalent in the stool samples at the other time points are outperformed by other *Streptococcus* species or strains at this stage in the gut microbiota development, before reaching a new bloom within 1 year of age. However, this possible temporal variation needs to be further investigated.

In summary, this comprehensive screening of samples from infant and maternal body sites reveals streptococcal species that are likely to thrive in the human gut environment. Further, distinct habitats, where the gut habitat was one of them, were observed for closely related species, possibly due to niche adaptations evolved in genes encoding proteins that facilitate host surface attachment. Further research, performed on a larger cohort and additional streptococcal isolates, is needed to determine phenotypic traits involved in gut microbiota colonization. Particularly, mechanistic studies on adhesins are essential to determine their contribution to body site specificity among closely related commensal streptococci.

## Materials and Methods

### Sample description

The 751 samples in this study were collected from 100 mother-infant pairs participating in the ProPACT study (41, 53, 54). The majority of the infants were born at term and breastfed during the first 3 months post-delivery (41). The infant stool samples were collected 10 days after birth (n = 100), 3 months after birth (n= 86), 1 year after birth (n = 89) and 2 years after birth (n = 88). The infant oral swabs (n = 72) were collected 10 days after birth. Stool samples (n = 93) and vaginal swabs (n = 96) from the mothers were collected in week 30 of pregnancy. In addition, 42 vaginal swabs were collected at birth. Breast milk samples (n = 85) were collected 10 days after birth. Out of the 100 mother-infant pairs, 20 pairs had a complete set of samples. The samples were stored at -40^°^C (vaginal swabs and breast milk) and -80^°^C (stool) before the analysis.

The study was approved by the Regional Committee for Medical Research Ethics Central Norway (Ref. 2012/2123).

### Isolation of *Streptococcus* from infant stool samples collected at 10 days of age

For selection of streptococci, Streptococcal Selective Agar C.O.B.A. plates (Thermo Scientific Oxoid) were used for cultivation. The 100 infant stool samples, stored in Cary-Blair transport medium (St. Olavs Hospital, Norway) at -80^°^C, were thawed on ice before 50 µl from each sample was spread onto C.O.B.A. plates. The plates were incubated anaerobically, using a GasPak system and AnaeroGen sachets (Oxoid, Thermo Scientific), at 37^°^C for 2 days. After the incubation, a maximum of 5 colonies with unique morphologies were picked from each plate and inoculated in 5 ml Todd Hewitt broth (Millipore), followed by an anaerobic incubation in the GasPak system at 37^°^C overnight. Glycerol stocks from each culture with visible growth were made and stored at -80^°^C. To identify the isolated bacteria, Sanger sequencing of the 16S rRNA gene was performed on amplicons generated with the cover all primer pair from Genetic Analysis (55).

### DNA extraction

DNA extraction of the infant oral swabs, the infant stool samples collected 3 months, 1 year and 2 years after birth, and the mothers’ stool samples were performed prior to this study (41, 43), using the LGC Mag Midi DNA extraction kit (LGC Genomics, UK). DNA extraction of the infant stool samples (10 days), vaginal samples and breast milk samples, as well as the streptococcal isolates and a ZymoBIOMICS Microbial Community Standard (Zymo Research, USA), was performed using the Zymo Research Quick-DNA Fecal/Soil Microbe 96 MagBead kit (Zymo Research, USA). The samples were extracted following the manufacturer’s recommendations, with some exceptions elaborated below. Before the extraction, the sample material was concentrated. Specifically, for the infant stool samples (10 days), 500 µl of sample was centrifuged at 13000 rpm for 5 min at 4^°^C, before the supernatant was removed. Similarly, 500 µl from the vaginal and breast milk samples were centrifuged at 21 500 x g for 30 min at 4^°^C before the supernatant was removed. The pellets from all the sample types were resuspended in 350 µl lysis buffer (Bashing Bead Buffer) and transferred to FastPrep tubes containing acid-washed glass beads of three sizes; 0.2 g of the <106 µm beads, 0.2 g of the 425-600 µm beads and 2 of the 2.5-3.5 mm beads (Sigma-Aldrich, Germany). *E. coli* 11775 was used as a positive DNA extraction control and lysis buffer (Bashing Bead Buffer) was used as a negative DNA extraction control. The samples were treated in a TissueLyser II (Qiagen) for 2 × 2.5 min at 30 Hz and centrifuged at 13000 x g for 1 min. The DNA was eluted in 30 µl elution buffer to ensure adequate DNA concentrations and quantified with the Qubit dsDNA HS assay kit (Thermo Fisher Scientific, USA).

### Library preparations for whole genome and metagenome sequencing

Library preparation of the stool samples (10 days) was performed with the Illumina DNA Prep kit (Illumina, USA), adding 2-250 ng DNA. Extracted DNA from the ZymoBIOMICS Microbial Community Standard and ZymoBIOMICS Microbial Community DNA Standard (Zymo Research, USA) was included. The pooled library was sequenced on NovaSeq6000 (Illumina, USA) at Novogene (UK), and 30 million reads per sample (2×150 paired-end reads) were requested. Library preparation of the streptococcal isolates, and the following sequencing on NovaSeq 6000 (Illumina, USA) (2×150 paired-end reads) was performed by Novogene (UK).

### Whole genome and metagenome data processing and analysis

The metagenome-assembled genomes (MAGs) were constructed as follows: The metagenomic raw reads were filtered and trimmed with BBDuk (https://sourceforge.net/projects/bbmap/) and mapped to the human genome using Bowtie 2 (56) for decontamination of human DNA. The decontaminated reads were assembled with SPAdes (57). The binning of assembled contigs was done twice using MaxBin2 (58) and MetaBAT2 (59). The MAGs were taxonomically classified using the newest release (R220) of the GTDB database (60).

Reconstruction of the isolate genomes included filtering, trimming, assembly of reads and taxonomic classification of assembled contigs, using the same tools and databases as mentioned for the MAGs. To establish the phylogenetic relationship between the isolates, estimated evolutionary distances were calculated using the MASH tool (61). The distances revealed that some of the isolates were phylogenetically identical. Based on this, a new assembly was done where the reads from the identical isolates were combined to increase coverage, resulting in 9 strain assemblies.

The phylogenetic relationship between the isolate assemblies, streptococcal MAGs and genomes in the GTDB database was estimated with MASH distances. From the GTDB database, all genomes from each of the isolate/MAG species were selected. Of these, only the genomes with the highest available assembly quality were used further. The GTDB genomes were clustered based on the MASH distances to find one representative genome for each species. A hierarchical clustering of these representative genomes, isolate assemblies and MAGs was done based on the MASH distances.

The strain assemblies of *S. parasanguinis I* and *S. parasanguinis F* were compared, aiming to find genetic differences. A blast database was made from the *S. parasanguinis I* assembly using BLAST+ (62). Then, the genes in the *S. parasanguinis F* assembly were identified with Prodigal (63) and blasted against the generated database using blastn. The same process was done the other way around, making a database from *S. parasanguinis* F and blasting the genes from *S. parasanguinis* I against it. The blast tables were filtered, keeping the hit with the best bit score in each case. Looking for dissimilar genes, the hits with alignment length to query length ratio lower than 0.75 were kept. The hit query sequences were translated to proteins, and a blastp search against the nr-database at NCBI (64) was done for identification. Protein sequence alignments between the matching genes were performed with Clustal Omega (1.2.4) (65, 66), and the protein domains were identified using InterProScan (67) with the InterPro database (68).

## Data availability

The whole genome sequencing data of the 16 streptococcal isolates are available in the NCBI SRA database, with the BioProject ID PRJNA1168056.

The 112 MAGs and their corresponding coverage table are available in https://arken.nmbu.no/~larssn/MiDiv/ida/.

## Primer design and testing

The nine streptococcal strain assemblies were used as template sequences, and the RUCS tool was used for primer generation (69). The specificity of a primer-pair was ensured by specifying for each chosen positive genome, a set of negative genomes where potential primers could not produce matches. For each assembly used as positive, the other assemblies, as well as all the MAGs from the infant stool samples, were set as negatives. In the cases where a MAG was very similar to an isolate, the MAG was set as positive as well.

The primers were tested *in silico* against all the strain assemblies and MAGs, using Geneious Prime 2023.1.2 (https://www.geneious.com), to validate their specificity. Similarly, potential binding towards common gut bacterial genomes from the NCBI Reference Sequence Database was investigated (Table S7). Two primer pairs for each strain were chosen based on their specificity and low hairpin melting temperature. These were finally tested with Primer-BLAST (70).

The specificities of the chosen primers were tested further using qPCR, by investigating their ability to amplify DNA from all the 16 streptococcal isolates. The annealing temperature of the primers was set to the same as their melting temperature (Tm = 60^°^C) to keep the occurrence of unspecific binding to a minimum. The reaction mix and amplification program used were the same as described for the streptococcal screening (see below). The best primer pair for each strain was determined by its specificity in the qPCR test run. The primer pair that showed the largest number of cycles between the true signal from the strain in question and signals from DNA amplification from other strains was used further. The characteristics of each primer can be found in Table S4.

### Screening of streptococcal strains by qPCR

The 751 samples were screened for the streptococcal strains. The total bacterial load in the samples was determined by amplification of the V3-V4 region of the 16S rRNA gene, using the primer pair PRK341F (5’-CCTACGGGRBGCASCAG-3’) and PRK806R (5’-GGACTACYVGGGTATCTAAT-3’) (71). The master mix for all qPCR runs contained 1x Hot FirePol EvaGreen qPCR Supermix (Solis Biodyne, Estonia) and 0.2 µM of each primer, in a reaction volume of 20 µl. In each run, DNA from the isolate in question was included as a positive control, and nuclease free water was used as a negative control. The amplification was carried out in a CFX96 Touch™ Real-Time PCR Detection System (Bio-Rad, USA). The initial denaturation step was executed at 95^°^C for 15 min, followed by 40 cycles of denaturation at 95^°^C for 30 sec and amplification and elongation at 60^°^C for 60 sec (streptococcal strain-specific primers) or amplification at 55^°^C for 30 sec and elongation at 75^°^C for 45 sec (16S rRNA gene primers). A melt curve analysis was included at the end of each run.

The qPCR results were analyzed with LinRegPCR (2014.x) (72) for baseline correction. To distinguish true qPCR signals from unspecific amplification, two criteria were set in the processing of the qPCR results; (1) the melting temperature had to match the melting temperature of the amplicon, and (2) the Cq value had to be two cycles lower than the negative control. These criteria are rather strict, lowering the streptococcal strain detection sensitivity, but ensuring a high specificity.

When a strain was detected in several samples from two different body sites, Fisher’s exact tests were used to test for associations between the samples from the different body sites, i.e. if there is a non-random co-occurrence of the strain in the two body sites. The p-values were corrected with a false discovery rate (FDR).

### Characterization of substrate utilization capability

The isolates were grown on different substrates in a Rapid ID 32 Strep test (bioMérieux, France). Prior to the test, isolate cultures were grown in Todd Hewitt broth (Millipore) and incubated anaerobically at 37 ^°^C overnight. The cultures were centrifuged to pellet the cells, which then were washed with 1X PBS and centrifuged once more. The cells were resuspended in MilliQ water to a turbidity of approximately McFarland standard 4. Further, the test was carried out according to the manufacturer’s recommendations, and the results were read manually after incubation for 4 hours and overnight. For each isolate, parallel tests were performed.

## Acknowledgments

We thank Ole Hjermundrud (master student, Norwegian University of Life Sciences) for contributing to the primer testing and Kaja Sundquist Wilhelmson (master student, Norwegian University of Life Sciences) for contributing to the qPCR screening.

This research was funded by the Faculty of Chemistry, Biotechnology and Food Science, Norwegian University of Life Sciences, NORWAY and the Research Council of Norway through the project UnveilMe: Unveiling the role of microbial metabolites in human infant development (project number 301364).

