## Supplemental_figures_and_tables for "Tracing of streptococcal strains from infant stool across human body sites links gut specificity to adhesins"

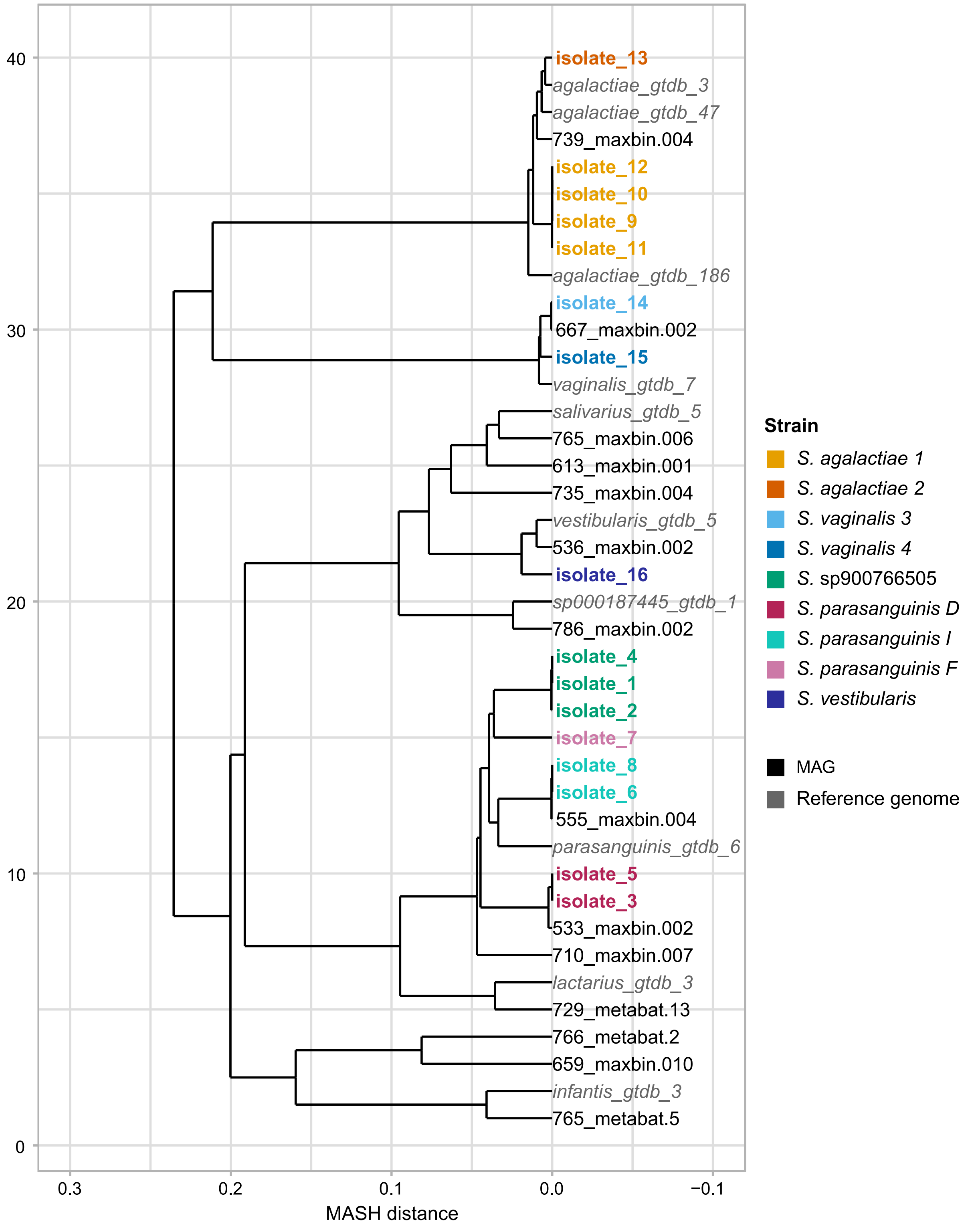


**Fig. S1. Phylogeny of streptococcal isolates and MAGs.** The mash distances between streptococcal isolates, streptococcal MAGs, and GTDB reference genomes are illustrated in a dendrogram. The isolates are colored according to their strain. The MAGs are shown in black and the GTDB reference genomes are marked in italic and grey.

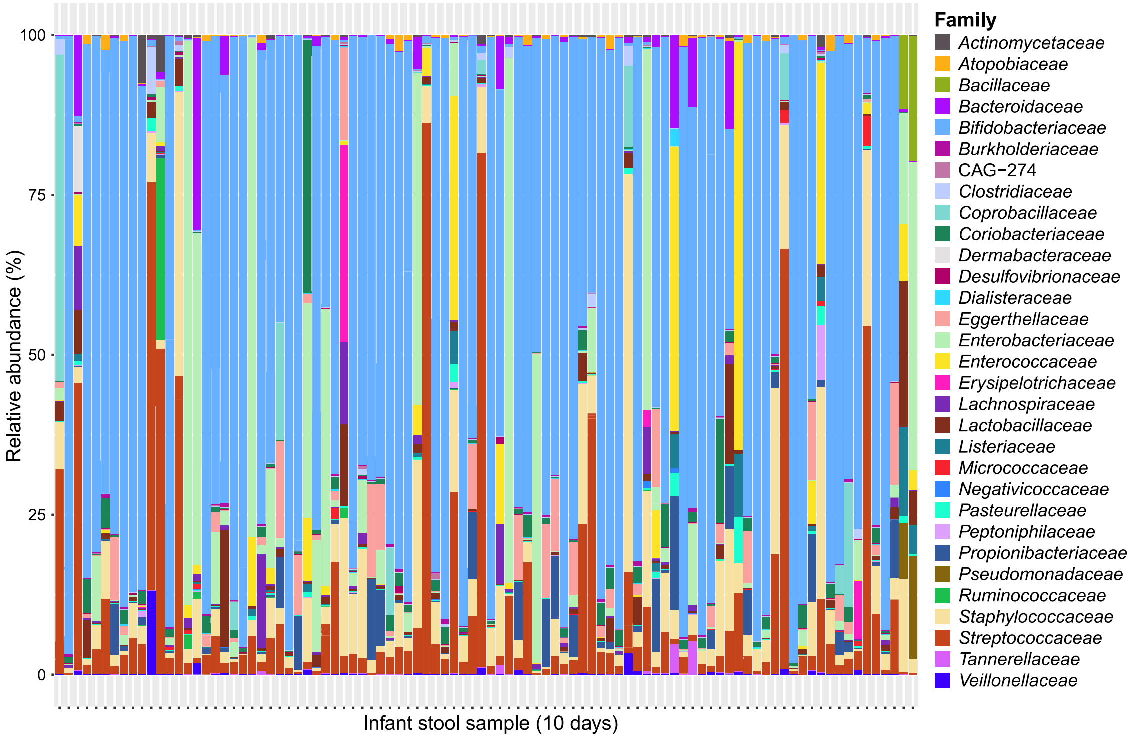


**Fig. S2. Bacterial composition in infant stool samples.** The figure shows the bacterial composition of the infant stool samples (10 days) at family level. Note that about half of the reads in the samples are not accounted for in this figure because they did not get assembled to MAGs.

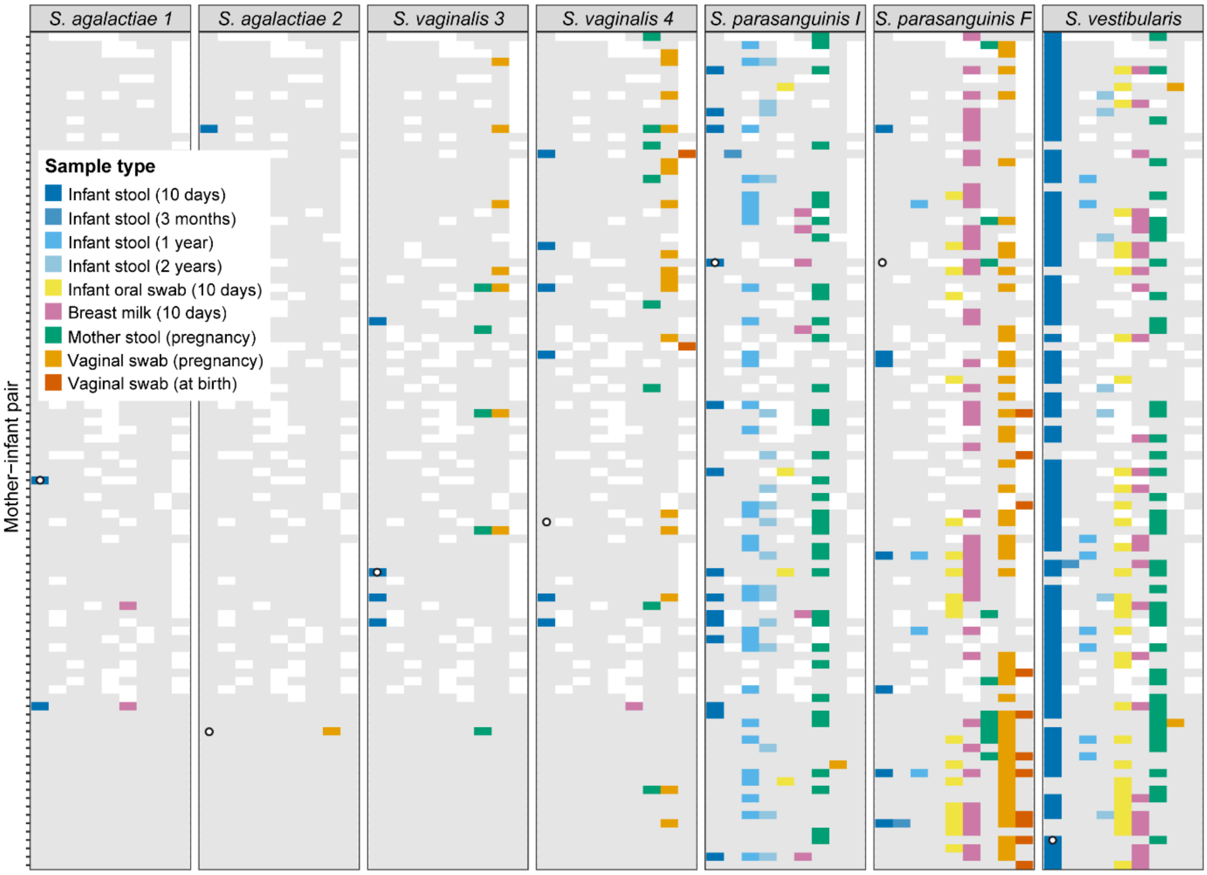


**Fig. S3. Streptococcal strain detections across different body sites.** The figure shows strain detections in infant stool collected at 10 days, 3 months, 1 year and 2 years of age (dark blue to light blue), infant oral swabs collected at 10 days of age (yellow), breast milk collected 10 days after birth (pink), mother's stool collected during pregnancy (green), vaginal swabs collected during pregnancy (light orange) and vaginal swabs collected at birth (dark orange). Samples where the strains are not detected are marked in grey, and missing samples are marked in white. The white circles indicate from which infant stool sample (10 days) the strains are isolated from.

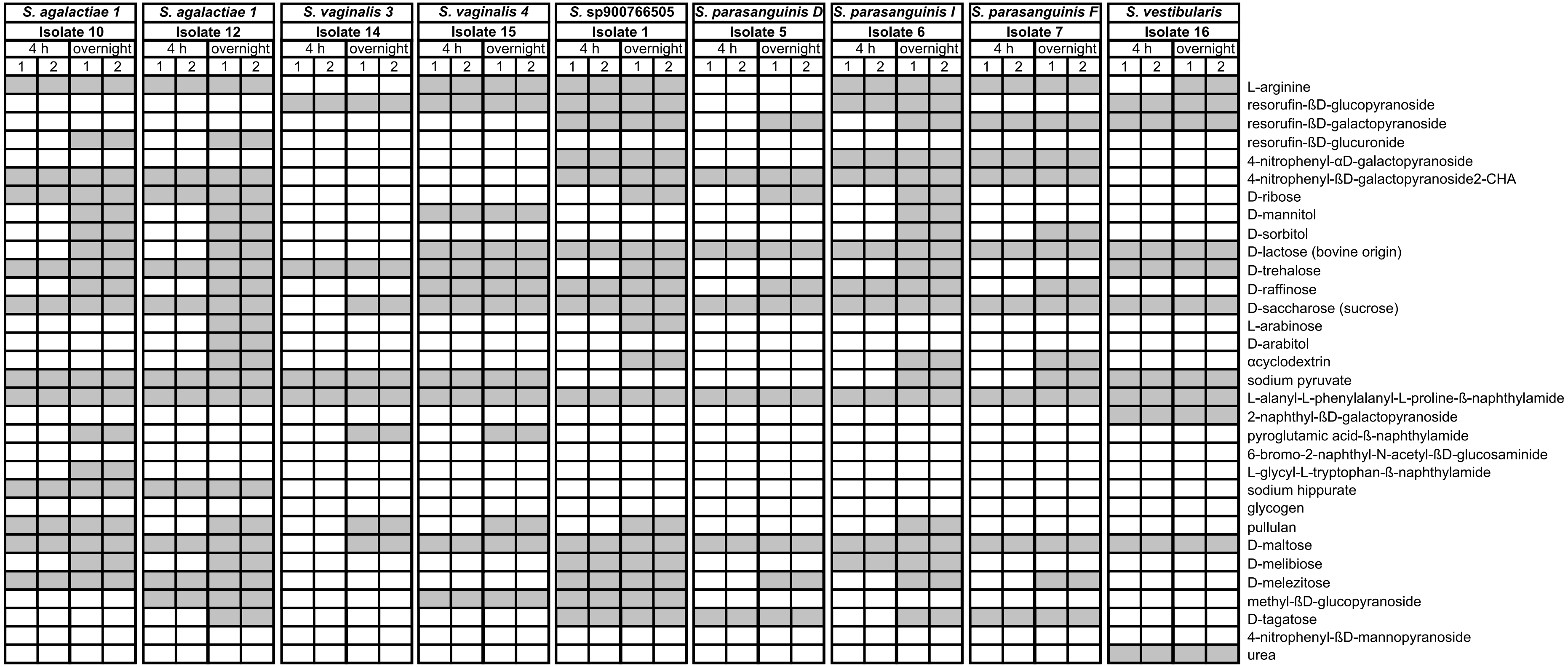


**Fig. S4. Substrate utilization.** The figure shows the strains capability to utilize different substrates, including a broad variety of carbohydrates. Positive reactions (grey) and negative reactions (white) after 4 hours of incubation and incubation overnight are shown. Parallel tests were performed for each isolate.

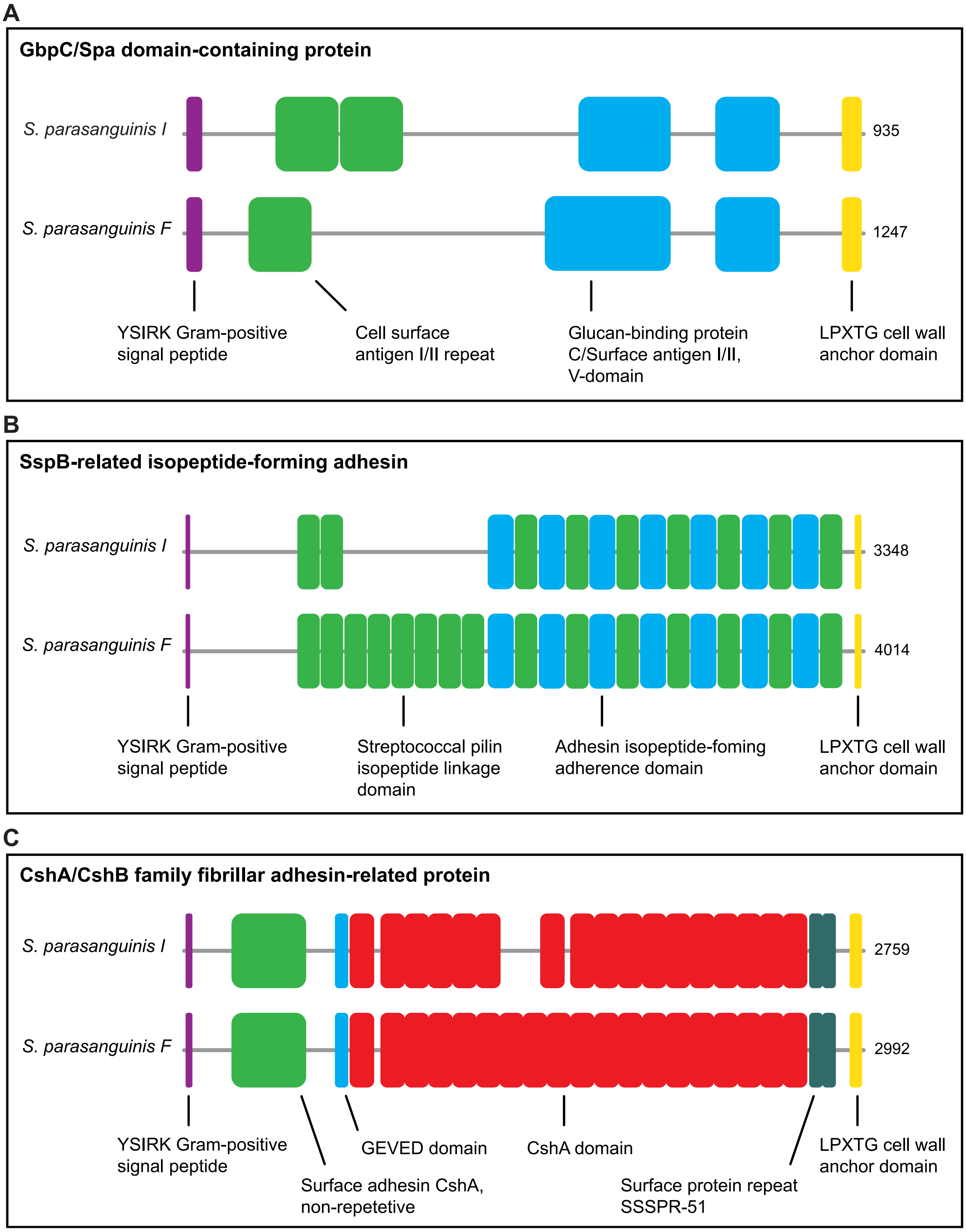


**Fig. S5. Illustrations of protein domains in adhesin-related proteins.** The figure shows *S. parasanguinis I and F* sequence alignments of protein domains in the GbpC/Spa domain-containing protein (A), the SspB-related isopeptide-forming adhesin (B) and the CshA/CshB family fibrillar adhesin-related protein (C). The colors of the domains correspond to the colors used to mark the same protein domains in the sequence alignments found in Table S6. The protein sequence length is noted at the end of each protein.

**Table S1. Taxonomic classification of the streptococcal isolates.** The table shows the GTDB database taxonomic classification of the streptococcal isolates.

| Isolate | Species | Strain names used in this work |
|---|---|---|
| 1 | *Streptococcus* sp900766505 | *Streptococcus* sp900766505 |
| 2 | *Streptococcus* sp900766505 | *Streptococcus* sp900766505 |
| 3 | *Streptococcus parasanguinis_D* | *Streptococcus parasanguinis D* |
| 4 | *Streptococcus* sp900766505 | *Streptococcus* sp900766505 |
| 5 | *Streptococcus parasanguinis_D* | *Streptococcus parasanguinis D* |
| 6 | *Streptococcus parasanguinis_I* | *Streptococcus parasanguinis I* |
| 7 | *Streptococcus parasanguinis_F* | *Streptococcus parasanguinis F* |
| 8 | *Streptococcus parasanguinis_I* | *Streptococcus parasanguinis I* |
| 9 | *Streptococcus agalactiae* | *Streptococcus agalactiae 1* |
| 10 | *Streptococcus agalactiae* | *Streptococcus agalactiae 1* |
| 11 | *Streptococcus agalactiae* | *Streptococcus agalactiae 1* |
| 12 | *Streptococcus agalactiae* | *Streptococcus agalactiae 1* |
| 13 | *Streptococcus agalactiae* | *Streptococcus agalactiae 2* |
| 14 | *Streptococcus vaginalis* | *Streptococcus vaginalis 3* |
| 15 | *Streptococcus vaginalis* | *Streptococcus vaginalis 4* |
| 16 | *Streptococcus vestibularis* | *Streptococcus vestibularis* |

**Table S2. Streptococcal MAGs.** The table shows the completeness, contamination and taxonomic classification of the streptococcal MAGs. The GTDB database was used for the taxonomic classification.

| **MAG id** | **Species** | **Completeness** | **Contamination** |
|---|---|---|---|
| 533_maxbin.002 | *Streptococcus parasanguinis_D* | 91.57 | 0.22 |
| 536_maxbin.002 | *Streptococcus vestibularis* | 100 | 0 |
| 555_maxbin.004 | *Streptococcus parasanguinis_S* | 100 | 0.01 |
| 613_maxbin.001 | *Streptococcus* sp001556435 | 100 | 0.11 |
| 659_maxbin.010 | *Streptococcus* spp. | 87.95 | 24.13 |
| 667_maxbin.002 | *Streptococcus vaginalis* | 100 | 0.17 |
| 710_maxbin.007 | *Streptococcus* sp900766505 | 82.37 | 1.86 |
| 729_metabat.13 | *Streptococcus lactarius* | 100 | 0.07 |
| 735_maxbin.004 | *Streptococcus* spp. | 77.76 | 22.25 |
| 739_maxbin.004 | *Streptococcus agalactiae* | 100 | 0.05 |
| 765_maxbin.006 | *Streptococcus salivarius* | 100 | 0 |
| 765_metabat.5 | *Streptococcus infantis_M* | 96.51 | 0.72 |
| 766_metabat.2 | *Streptococcus* spp. | 81.46 | 0.04 |
| 786_maxbin.002 | *Streptococcus* sp000187445 | 100 | 0.06 |

**Table S4. Fisher's test.** The table below shows the p-values, adjusted p-values (FDR) and odds ratios for the tested associations between strain detections in samples collected from mothers and infants.

| Isolate | Contingency table | | | p-value | p-value (adjusted, FDR) | Odds ratio |
|---|---|---|---|---|---|---|
| ***S. vestibularis*** | | **breast milk (10 days)** | | 6.21E-05 | 0.0025 | 10.7539 |
| | **oral swab (10 days)** | Detected | Not detected | | | |
| | Detected | 18 | 12 | | | |
| | Not detected | 4 | 30 | | | |
| ***S. vestibularis*** | | **mother stool (pregnancy)** | | 0.1053 | 0.7702 | 2.7464 |
| | **infant stool (10 days)** | Detected | Not detected | | | |
| | Detected | 18 | 12 | | | |
| | Not detected | 4 | 30 | | | |
| ***S. vestibularis*** | | **breast milk (10 days)** | | 0.3683 | 0.9118 | 2.0706 |
| | **infant stool (10 days)** | Detected | Not detected | | | |
| | Detected | 18 | 12 | | | |
| | Not detected | 4 | 30 | | | |
| ***S. vestibularis*** | | **mother stool (pregnancy)** | | 0.1429 | 0.7702 | 0.4565 |
| | **oral swab (10 days)** | Detected | Not detected | | | |
| | Detected | 10 | 19 | | | |
| | Not detected | 21 | 18 | | | |
| ***S. vestibularis*** | | **oral swab (10 days)** | | 1 | 1 | 1.1433 |
| | **infant stool (10 days)** | Detected | Not detected | | | |
| | Detected | 27 | 33 | | | |
| | Not detected | 5 | 7 | | | |
| ***S. vestibularis*** | | **mother stool (pregnancy)** | | 0.4666 | 0.9118 | 0.6311 |
| | **breast milk (10 days)** | Detected | Not detected | | | |
| | Detected | 9 | 16 | | | |
| | Not detected | 26 | 29 | | | |
| ***S. vestibularis*** | | **infant stool (1 year)** | | 1 | 1 | 1.2149 |
| | **infant stool (10 days)** | Detected | Not detected | | | |
| | Detected | 7 | 69 | | | |
| | Not detected | 1 | 12 | | | |
| ***S. vestibularis*** | | **infant stool (2 years)** | | 1 | 1 | 1.2507 |
| | **infant stool (10 days)** | Detected | Not detected | | | |
| | Detected | 6 | 67 | | | |
| | Not detected | 1 | 14 | | | |
| ***S. vestibularis*** | | **infant stool (2 years)** | | 1 | 1 | 0 |
| | **infant stool (1 year)** | Detected | Not detected | | | |
| | Detected | 0 | 8 | | | |
| | Not detected | 5 | 65 | | | |
| ***S. vestibularis*** | | **oral swab (10 days)** | | 0.69 | 1 | 0.5085 |
| | **infant stool (1 year)** | Detected | Not detected | | | |
| | Detected | 2 | 5 | | | |
| | Not detected | 27 | 34 | | | |
| ***S. vestibularis*** | | **breast milk (10 days)** | | 0.4147 | 0.9118 | 0.3164 |
| | **infant stool (1 year)** | Detected | Not detected | | | |
| | Detected | 1 | 6 | | | |
| | Not detected | 24 | 45 | | | |
| ***S. vestibularis*** | | **mother stool (pregnancy)** | | 0.449 | 0.9118 | 0.4613 |
| | **infant stool (1 year)** | Detected | Not detected | | | |
| | Detected | 2 | 5 | | | |
| | Not detected | 35 | 40 | | | |

| | | | | | | |
|---|---|---|---|---|---|---|
| ***S. vestibularis*** | | **oral swab (10 days)** | | 1 | 1 | 1.4154 |
| | **infant stool (2 years)** | Detected | Not detected | | | |
| | Detected | 2 | 2 | | | |
| | Not detected | 26 | 37 | | | |
| ***S. vestibularis*** | | **breast milk (10 days)** | | 0.4207 | 0.9118 | 0.3374 |
| | **infant stool (2 years)** | Detected | Not detected | | | |
| | Detected | 1 | 6 | | | |
| | Not detected | 22 | 44 | | | |
| ***S. vestibularis*** | | **mother stool (pregnancy)** | | 0.6937 | 1 | 0.5413 |
| | **infant stool (2 years)** | Detected | Not detected | | | |
| | Detected | 2 | 5 | | | |
| | Not detected | 32 | 43 | | | |
| ***S. parasanguinis F*** | | **breast milk (10 days)** | | 0.2574 | 0.7702 | 2.2407 |
| | **oral swab (10 days)** | Detected | Not detected | | | |
| | Detected | 11 | 6 | | | |
| | Not detected | 21 | 26 | | | |
| ***S. parasanguinis F*** | | **vaginal swab (pregnancy)** | | 0.8249 | 1 | 0.8237 |
| | **breast milk (10 days)** | Detected | Not detected | | | |
| | Detected | 21 | 23 | | | |
| | Not detected | 20 | 18 | | | |
| ***S. parasanguinis F*** | | **breast milk (10 days)** | | 0.2068 | 0.7702 | 4.7982 |
| | **infant stool (10 days)** | Detected | Not detected | | | |
| | Detected | 5 | 1 | | | |
| | Not detected | 40 | 39 | | | |
| ***S. parasanguinis F*** | | **vaginal swab (pregnancy)** | | 0.4361 | 0.9118 | 2.5332 |
| | **infant stool (10 days)** | Detected | Not detected | | | |
| | Detected | 5 | 2 | | | |
| | Not detected | 44 | 45 | | | |
| ***S. parasanguinis F*** | | **vaginal swab (at birth)** | | 0.1629 | 0.7702 | 6.2970 |
| | **infant stool (10 days)** | Detected | Not detected | | | |
| | Detected | 2 | 1 | | | |
| | Not detected | 9 | 30 | | | |
| ***S. parasanguinis F*** | | **vaginal swab (pregnancy)** | | 0.2625 | 0.7702 | 3.1597 |
| | **vaginal swab (at birth)** | Detected | Not detected | | | |
| | Detected | 8 | 2 | | | |
| | Not detected | 16 | 13 | | | |
| ***S. parasanguinis F*** | | **infant stool (1 year)** | | 0.02975 | 0.4073 | 14.8048 |
| | **infant stool (10 days)** | Detected | Not detected | | | |
| | Detected | 2 | 5 | | | |
| | Not detected | 2 | 80 | | | |
| ***S. parasanguinis F*** | | **oral swab (10 days)** | | 1 | 1 | 0.9227 |
| | **infant stool (1 year)** | Detected | Not detected | | | |
| | Detected | 1 | 3 | | | |
| | Not detected | 17 | 47 | | | |
| ***S. parasanguinis F*** | | **breast milk (10 days)** | | 0.1156 | 0.7702 | Inf |
| | **infant stool (1 year)** | Detected | Not detected | | | |
| | Detected | 4 | 0 | | | |
| | Not detected | 35 | 37 | | | |
| ***S. parasanguinis F*** | | **mother stool (pregnancy)** | | 1 | 1 | 0 |
| | **infant stool (1 year)** | Detected | Not detected | | | |
| | Detected | 0 | 3 | | | |
| | Not detected | 9 | 70 | | | |
| ***S. parasanguinis F*** | | **vaginal swab (pregnancy)** | | 1 | 1 | 0.9529 |
| | **infant stool (1 year)** | Detected | Not detected | | | |
| | Detected | 2 | 2 | | | |
| | Not detected | 42 | 40 | | | |

| | | | | | | |
|---|---|---|---|---|---|---|
| ***S. parasanguinis F*** | | **vaginal swab (at birth)** | | 0.2632 | 0.7702 | Inf |
| | **infant stool (1 year)** | Detected | Not detected | | | |
| | Detected | 1 | 0 | | | |
| | Not detected | 9 | 28 | | | |
| ***S. parasanguinis I*** | | **mother stool (pregnancy)** | | 0.0434 | 0.4449 | 6.6519 |
| | **infant stool (10 days)** | Detected | Not detected | | | |
| | Detected | 3 | 10 | | | |
| | Not detected | 3 | 69 | | | |
| ***S. parasanguinis I*** | | **mother stool (pregnancy)** | | 0.658 | 1 | 0.4750 |
| | **breast milk (10 days)** | Detected | Not detected | | | |
| | Detected | 1 | 4 | | | |
| | Not detected | 26 | 49 | | | |
| ***S. parasanguinis I*** | | **infant stool (1 year)** | | 1 | 1 | 1.0465 |
| | **infant stool (10 days)** | Detected | Not detected | | | |
| | Detected | 5 | 9 | | | |
| | Not detected | 26 | 49 | | | |
| ***S. parasanguinis I*** | | **infant stool (2 years)** | | 0.2123 | 0.7702 | 2.7722 |
| | **infant stool (10 days)** | Detected | Not detected | | | |
| | Detected | 4 | 7 | | | |
| | Not detected | 13 | 64 | | | |
| ***S. parasanguinis I*** | | **infant stool (2 years)** | | 0.258 | 0.7702 | 1.9349 |
| | **infant stool (1 year)** | Detected | Not detected | | | |
| | Detected | 8 | 21 | | | |
| | Not detected | 8 | 41 | | | |
| ***S. parasanguinis I*** | | **oral swab (10 days)** | | 1 | 1 | 0.7921 |
| | **infant stool (1 year)** | Detected | Not detected | | | |
| | Detected | 1 | 19 | | | |
| | Not detected | 3 | 45 | | | |
| ***S. parasanguinis I*** | | **breast milk (10 days)** | | 1 | 1 | 0.9012 |
| | **infant stool (1 year)** | Detected | Not detected | | | |
| | Detected | 2 | 25 | | | |
| | Not detected | 4 | 45 | | | |
| ***S. parasanguinis I*** | | **mother stool (pregnancy)** | | 0.8095 | 1 | 1.2004 |
| | **infant stool (1 year)** | Detected | Not detected | | | |
| | Detected | 11 | 16 | | | |
| | Not detected | 20 | 35 | | | |
| ***S. parasanguinis I*** | | **oral swab (10 days)** | | 1 | 1 | 0 |
| | **infant stool (2 years)** | Detected | Not detected | | | |
| | Detected | 0 | 15 | | | |
| | Not detected | 3 | 49 | | | |
| ***S. parasanguinis I*** | | **breast milk (10 days)** | | 1 | 1 | 1.0569 |
| | **infant stool (2 years)** | Detected | Not detected | | | |
| | Detected | 1 | 13 | | | |
| | Not detected | 4 | 55 | | | |
| ***S. parasanguinis I*** | | **mother stool (pregnancy)** | | 1 | 1 | 0.9842 |
| | **infant stool (2 years)** | Detected | Not detected | | | |
| | Detected | 6 | 10 | | | |
| | Not detected | 25 | 41 | | | |
| ***S. vaginalis 4*** | | **vaginal swab (pregnancy)** | | 0.2869 | 0.7845 | 2.4700 |
| | **infant stool (10 days)** | Detected | Not detected | | | |
| | Detected | 2 | 4 | | | |
| | Not detected | 15 | 75 | | | |
| ***S. vaginalis 4*** | | **vaginal swab (pregnancy)** | | 0.6309 | 1 | 1.5858 |
| | **mother stool (pregnancy)** | Detected | Not detected | | | |
| | Detected | 2 | 6 | | | |
| | Not detected | 14 | 67 | | | |

| | | | | | | |
|---|---|---|---|---|---|---|
| ***S. vaginalis 3*** | | **vaginal swab (pregnancy)** | | 0.00287 | 0.0588 | 26.8614 |
| | **mother stool (pregnancy)** | Detected | Not detected | | | |
| | Detected | 3 | 2 | | | |
| | Not detected | 4 | 80 | | | |

**Table S7. Gut bacterial genomes used in primer testing.** The gut bacterial genomes listed in the table were downloaded from the NCBI Reference Sequence Database and used for testing the specificity of the designed primers.

| NCBI RefSeq assembly / Accession | Species |
|---|---|
| GCF_000008865.2 | *Escherichia coli* O157:H7 |
| GCF_000146185.1 | *[Eubacterium] eligens* |
| GCF_000154385.1 | *Faecalibacterium prausnitzii* |
| GCF_000393015.1 | *Enterococcus faecalis* |
| GCF_000439915.2 | *Lactobacillus gasseri* |
| GCF_002891025.1 | *Enterococcus avium* |
| GCF_006094375.1 | *Staphylococcus epidermidis* |
| GCF_009734005.1 | *Enterococcus faecium* |
| GCF_014131755.1 | *Bacteroides thetaiotaomicron* |
| GCF_018292205.1 | *Bacteroides caccae* |
| GCF_020541885.1 | *Bifidobacterium pseudocatenulatum* |
| GCF_024397795.1 | *Lactobacillus intestinalis* |
| GCF_032486935.1 | *Winkia neuii* |
| NC_000913.3 | *Escherichia coli* str. K-12 substr. MG1655 |
| NC_007795.1 | *Staphylococcus aureus* subsp. *aureus* NCTC 8325 chromosome |
| NC_015067.1 | *Bifidobacterium longum* subsp. *longum* JCM 1217 |
| NZ_AKCA01000001.1 | *Bifidobacterium bifidum* NCIMB 41171 cont1.1 |
| NZ_CP014022.1 | *Staphylococcus lugdunensis* strain FDAARGOS_141 chromosome |
| NZ_CP040780.1 | *Lacticaseibacillus rhamnosus* strain 1.0320 chromosome |
| NZ_CP068170.1 | *Thomasclavelia ramosa* strain FDAARGOS_1105 chromosome |
