## Supplemental Table S6 for "Tracing of streptococcal strains from infant stool across human body sites links gut specificity to adhesins"

| | |
|---|---|
| **Protein:** | GbpC/Spa domain-containing protein |
| **Gene location in *S. parasanguinis I* genome:** | NODE_10_length_105029_cov_696.487737;11638;14445;- |
| **Gene location in *S. parasanguinis F* genome:** | NODE_8_length_96838_cov_337.469879;81419;85162;+ |

| Color code | Protein name | Location (para_I) | Location (para_F) |
|---|---|---|---|
| Color | YSIRK Gram-positive signal peptide | 10-29 | 10-29 |
| Color | Cell surface antigen I/II A repeat | 157-234 / 238-314 | 115-192 |
| Color | Glucan-binding protein C/Surface antigen I/II, V-domain | 386-554 / 638-739 | 633-859 / 944-1044 |
| Color | Glucan-binding protein C/Surface antigen I/II, V-domain superfamily | 284-539 / 574-748 | 594-844 / 944-1060 |
| Color | LPXTG cell wall anchor domain | 898-934 | 1210-1246 |

```
para_I MRSYREHVSKQEKFSIRKLSVGVVSLAIAGLATVNTYGAEVKADEATAPSTEATSTDTAT      60
para_F MKSYREHVSKQEKFSIRKLSVGVVSLAIAGLATVNTYGAEVKADETTAPATEATTTDTAT      60
       *:*******************************************:***:****:*****

para_I SDAAVATSSKLTSTTVTEGNKIVTTTYVESPELEKAKADAATEGVTVTEEAEKVQPSIVA      120
para_F SDAAVATSSKLTSTSVTEGNKTVTTTYVESPELEKAKADAATEGVTVTEEAEKIQPSIAA      120
       **************:****** *******************************:****.*

para_I AEADNKAQTAEINTVVENYKKAKAEYEAKSQEITLIEKRNAEAEAAYKKQVEDYNSQQA-      179
para_F AEADNKAQTAEINTVVENYKKAKAEYEAKLAKQKQVEAENAKAKADYDQANTEYQNDLLA      180
       *****************************  : . :* .**:*:* *.:   :*:.:

para_I ------------------------------------------------------------      179
para_F YQAQKAEYEKAKASYAESQKAYEEALAAYKAASTSTSSKDSDSYKLVKEQYEAAKATYET      240

para_I -------------------------------AYKAALAAYNQKKVAYDA-----------      197
para_F SKAKYETDAAKYADNELSFTDATASYNTAKTSYDAALAQYNVAKAQYDKELESYESKKAE      300
                                      :*.**** **   *. **

para_I ---------------------------------------------KLAEKAAA-------      205
para_F FEKNKLAQAAAENKFDQQTKEYTAAEKQYQQDLAAYNEAKTAYDTALVAYNVATSTGGTD      360
                                                     *.   .*

para_I DKANAEAKAKYEAEMAVYNTAKAQYDKDLQEYQAKKAQYDKDKEAYGKLVA----KKAEE      261
para_F SAEYAALKAKYEAEKSAYETAKAKYDVDAENYNDKKTSYETAKASYDSTKTAYDVAKSSY      420
       .   *  ******* :.*:****:** * ::*: **:.*:. * :*..  :     *:.

para_I DAAKAKYEVELAKYNVDLEQ----------------------------------------      281
para_F DIAKTQYDSEMAKYNKATEEYNKAVSDYQKAQEQYAKDKLEYDTAATSYAKAKTEYDTVV      480
       * **::*: *:****   *:

para_I ------------------------------------------------------------      281
para_F NKFNKFEKEIKAIYKNATVNIVESNTIEEDEEKEVEKDKDGKPTKELTKKAKDLTAAKNL      540
```

```
para_I -----------------------------------YG-----------KDFATYQTKLAE     295
para_F YKNTKSQYEEDKKAFEEAKAKYASNEIEFVAVKAAYDKAKSSYDAVNEQYAKLYQAKLAS     600
                                          *.           :    **:***.

para_I YQTALAQYKDAKAAYDKYMNDNGFQDLKDVETVQDLTFQREGGATHTIDGISTYLTRDAQ     355
para_F IETALAQYKNAKEAYEKYMTDNGFKDLKDVETVQDLTFQREGGATHTIDGISTYLTRDAQ     660
        :*******:** **:***.****:***********************************

para_I ARLNTSNVHQYDSNKLEASDIVATSPWANNETEFIQIKEGDKFVVTYDNLNQSSMRENKK     415
para_F KRLNKSNVHQYDSNELEASDIVATSPWTNNETEYIQVKEGDKFVVTYDNLNQSSMREN-T     719
        ***.*********:************:*****:**:********************* .

para_I DMHPIKRVIYRYEILSLPSNDGKGIAAISADPTVTMTVGAST-DQNKPVKVAVDVEFYDG     474
para_F DMHPIKRVIYRYEILSLPSKDGKGIAAVSNDPTVTLTVGASTDDQDKPVKVAVDVEFYDG     779
       *******************:*******:* *****:****** **:**************

para_I DGNKFDLTQHNAIVALNSLNHWTGASYVDSGDKPRALTVEAKDKNGNTVRGTWNPYADGS     534
para_F NGNKFDLTQHNAIVALNSLNHWTGASYVDSGDKPRTLTVEAKDKNGNTVRGTWNPYADGS     839
       :**********************************:************************

para_I SMSIENNAVVVKNGTADFGTADVTISAENPIKIVAQNATWNGSEFAVSEETVIDATSVNA     594
para_F SMSIENNAVVVKNGKADFGTADVTISAENPIKIVAQKATWNGSEFAVSEETVIDATSVNT     899
       **************.*********************:**********************:

para_I SGAGNGHDIGTDEFTLDGKDDVIGSYTIDPTSGRIIFTPKKKFENVEHQESVNVGNNKYI     654
para_F SGGGNGHSIGTEEFTFEGKDDVLGSYKVDATTGQITFTPKKKFENVEHQESVNIGNNKYI     959
       **.****.***:***::*****:***.:* *:*:* *****************:******

para_I PIPNSSVTYDSATKEVTSFKDNQYIEHGSIFNGESSTTLEGWDNPSSPYLYYGGAGLKMS     714
para_F KIPNSSVSYDSATKEVTSERDNQYVEHGAVFNGESSATLRGWDDPSSPYLYYGGAGLKMS     1019
        ******:********** :****:***::******:**.***:****************

para_I DGHLVFTANGANAAGQPTVYWFAINSNVGIPKNPGEAPKEPTKPTEPKAPTPPTITVENL     774
para_F DGHLVFTAKGANAAGQPTVYWFAINSNVGLPKNPGSFVIPP-APVAPVAPVEPKLNEEK-     1077
       ********:********************:*****.    *  *. * **. *.:. *:

para_I PAEPTKPEEPKSPTPPTAPNYTVI----------TVDVEEPKAPTPPKAPTPPTPEVVPN     824
para_F PTLP-TPLNPVSPVAPVKPVKEEIEKIKEPTPPTPVAPVSPVSPIEPKAPEQPKYKTIPK     1136
       *: * .* :* **. *. *    *           *   .* :*  ****  *. :.:*:

para_I PVKPEKAEVKWHKNKVVTETDIPTPPTPVPPTPPTPYNPPTPIVPPTPIVPPTPEVPTEP     884
para_F TEEPQSAEVKWHKNKVVTETDIPTPPTPVPPTPPTPYNPPTPIVPPTPIVPPTPEVPTEP     1196
         :*:.******************************************************

para_I TPEVPEQPVQPAQPQTPALPNTGTESSAAAVLAGAMAGLLGLGLARKKKED  935
para_F TPEVPEQPVQPAQPQTPALPNTGTESSTAAVLAGAMAGLLGLGLARKKKED  1247
       ***************************:***********************
```

| | |
|---|---|
| **Protein:** | accessory Sec-dependent serine-rich glycoprotein adhesin |
| **Gene location in *S. parasanguinis I* genome:** | NODE_12_length_79046_cov_686.224962;65370;70337;- |
| **Gene location in *S. parasanguinis F* genome:** | NODE_16_length_50571_cov_308.191013;1280;5845;+ |

| **Color code** | **Protein name** | **Location (para_I)** | **Location (para_F)** |
|---|---|---|---|
| Color | KxYKxGKxW signal peptide | 14-45 | 14-45 |
| Color | Serine-rich repeat adhesion glycoprotein, N-terminal domain | 36-84 | 36-84 |
| Color | Cell Division and Developmental Signaling Domain-Containing Protein | 316-1027 | |
| Color | LPXTG cell wall anchor domain | 1621-1655 | 1487-1521 |
| Color | Overlapping domains | | |

```
para_I MQFKRSKGNFRETDRVVRFKLIKSGKNWLRASTAALGLFRVVRGQVEETIIANVQQDQIE      60
para_F MQFKRSKGNFRETDRVVRFKLIKSGKNWLRASTAALGLFRVVRGQVEETIIANVQQDQIE      60
       ************************************************************

para_I NQKHNQAFLKGLITVGTVFGGAVLATTAKAEDATSLAPTVSETKEETLAEVDSVVLGNTS      120
para_F SQKHNQAFLKGLITVGTVFGGAVLATTAKAEDATSLAPTVSETKEETLAEVDSVVLAKTS      120
       .*******************************************************.:**

para_I TQSSESSSVSGSTSLSTSVSVSTSISESASLSLSEVGSTALSTALSESAQALESEVVVDE      180
para_F NQPSESISVSESASFSTSESASASISESTSLSLSEVGSTALSTALSESAQALESEVVVEE      180
       .* *** *** *:*:*** *.*:*****:*****************************:*

para_I PTSLEEATVLEQNTSEAELLQEIAGNYASKMTDNDRRSVVEAVINKVQAEVTASNNLIHT      240
para_F PTSLEEAVVLEQNTSEAELLQEIAGNYASKMTDTDRRAVVQAVINKVQAEVTASNNLIHT      240
       *******.*************************.***:**:*******************

para_I NASAQAYADQRDRLEKAVDEMMTTLTAAGFVGNTNVDGKPAISAQLAPLAEETYLADDVL      300
para_F NASAQAYADQRDRLEKAVDEMMTTLTAAGFVGNTNVDGKPAISAQLAPIAEETYLADDVL      300
       ************************************************:***********

para_I DMSPNPEDPNGASVEDPTLDTPGYAKDPHLDKDLLRFSPEELKNFLGEPYTNRYTFGIWD      360
para_F DMTPNPEDPNGASVEDPTLDTPGYAKDPHLDKDLLRFSPEELKNFLGEPYTNRYTFGIWD      360
       **:*********************************************************

para_I FVNKQGESLGYYATMSIDISEIDPEKHKAMDVYFRIVRKSDGAEIFSQTVSPKGYGQDIQ      420
para_F FVNKQGESLGYYATMSIDISEIDPEKHKAMDVYFRIVRKSDGVEIFSQTVSPEGYGQDIQ      420
       ******************************************.*********:*******

para_I LPKEVLIGQAPFGNKVFNSEPSTDNGTFGMMTNFIPEQAFLFRSIYDVMTPENQGQLIRT      480
para_F LPKEVLIGQAPFGNKVFNSKPTTGNGTFGMLTNFIPEQAFLFRSIYDVMTPENQGQLIRT      480
       *******************:*:*.******:*****************************

para_I YPSIKIPSMMGQQSTFYREVDPNGRWFNGQYEPTGKERSLLEYRIYGLEGQHYTASNPRE      540
para_F YPSIKIPSMMGQQSTFYREVDPNGRWFNGQYEPTGKERSLLEYRIYGLEGQHYTASNPRE      540
       ************************************************************

para_I FPGYVQVPAHTVFPNRKSGVFDNSKNGKSRIELLGDAREHFIKSEVVTLNQNGDYSLRYY      600
para_F FPGYVQVPAHTVFPNRQSGVFDNSKNGKSRVELLGDAREHFIKSEVVTLNQNGDYSLRYY      600
       ****************:*************:*****************************
```

```
para_I VLDPSKIHDVSSGDVGNTQVTDVYTLVYEKEFKQDSTDKLSDLGGTRKVESKNKDYFLNV      660
para_F VLDPSKIHDVSSGDVGNTQVTDVYTLVYEKEFKQDSTDKLSDLGGTRKVESKNKDYFLNV      660
       ************************************************************

para_I TPRRIDYQHFELDITGWFSSKETATYTDEKTGIEYTVPKPFTELPKSAYLDKNTTMVVGE      720
para_F TPRRIDYQHFELDITGWFSSKETVTYTDEKTGIEYTVPKPFTELPKSAYLDKNTTMVVGE      720
       ***********************.************************************

para_I DATPQGADGFSNFKQTIKKIVDSYPLTSVNYYYRKMTPSESASQSQNFSISTSESLVSES      780
para_F DATPQGADGFSNFKQTIKKIVDSYPLTSVNYYYRKMTPSESASQSQNFSILTSESLVSES      780
       ************************************************** *********

para_I ISSSQSVSVSESVSLSQASSSASQSDSQVSTSISLSQSVSTSESNSLVQESVSASQSESL      840
para_F ISSSQSVSVSESVSLSQASSSASQSDSQVSTSISLSQSVSTSESNSLVQESISTSQSEAL      840
       ***************************************************:*:****:*

para_I VQESVSASQSESLVQASVSASQSDSFSASQSDSLVQASVSSSQSESLVQASVLASESNSL      900
para_F VQESVSASQSDSLVQSSVSASQSE--------SLIQASVSASQSD-------------SL      879
       **********:****:*******:        **:*****:***:             **

para_I AQESISASQSDSLVQASVSASQSNSFVQALVSASQSESLLQESVSASQSDSLVQASVSSS      960
para_F VKESVSASQSDSLVQASVSASQSDSLVQSSVSASQSESLIQASVSASQSESLVQDSVSAS      939
       .:**:******************:*:**: *********:* *******:**** ***:*

para_I QSDSLVQASVSASQSESLVQASVSASQSESLVQASVSASQSESLVQASVSVSQSQSTSDV      1020
para_F QSDSLIQESVSASQSDSLIQSSVSSSQSESLVQASVSASRSESLVQESVSAS--------      991
       *****:* *******:**:*:***:**************:****** ***.*

para_I ESKSQITESMSFSRSDSMSQSETQVSTDLQSTSLSQSESLVRESVSASESESLVQESVSA      1080
para_F -----------------------------------QSDSLIQESVSASQSDSLVQASVSA      1016
                                          **:**::******:*:**** ****

para_I SQSDSLIQESVSASQSESLVQESVSASQSESLTQASVSASQSESLVQESVSASESELLVQ      1140
para_F SQSESLVKESVSASQSESLLQESVSASQSDSLNQASVSASRSESLVQASVSASQSESLVQ      1076
       ***:**::***********:*********:**.*******:****** *****:** ***

para_I ESVSASQSDSLIQESVSASQSDSLVQESISASQSESLIQASVSESQSQSTSDIESKSQIT      1200
para_F ESVSVSQSDSLIQASVSASQSDSLVQVSVSASQSESLVQASVSAS---------------      1121
       ****.******** ************ *:********:***** *

para_I ESMSFSRSDSMSQSESQVSTDLQSTSLSQSESLVRESVSASESESLVRESVSASQSESQV      1260
para_F -----------------------------------------QSDSLVQASVSASQSESLL      1140
                                                :*:***: ********* :

para_I QVSVSASQSDSLVQASVLTSQSESLVQESISASQSDSLVQESVSTIQSDSLAQASVSTSQ      1320
para_F QESVSASQSDSLVQESVSSSQSESLVQESISASQSESLIKESVSASQSDSQSKAASASKS      1200
       * ************ ** :****************:**::****: **** ::*: ::..

para_I SESLIQDSVSASQSDSLVQGSISASQSESLVQESVSASQSDSLAQASVSASQSDSLVQES      1380
para_F ESEKASKSVSLSQSVSL----------SNLVSRSISVSQSQST-----SAVGSEEVVSQL      1245
       ...  ..*** *** **          ..**..*:*.***:*      **  *:.:*.:

para_I VSASQ---SDSLIQASVSASQSESLVQESVSSSQSDSLAQASVSASESNSLAQASVSASQ      1437
para_F ISLSQSISMSEKLSESVSMSQSESLVQASVSASQSDSLVQASISASQSESQVQASVSASQ      1305
       :* **    .. :. *** ******** ***:******.***:***:*:* .********

para_I SDSLVQASVSASQSDSLVQESVSASQLDSLVQASASTSESASTSHVVAASQVSESGSFSR      1497
para_F SESLIQESILANQSESLVQASISSSQSDSLVQASASTSESASTSHVVAASQVSESGSFSR      1365
       *:**:* *: *.**:**** *:*:** *********************************
```

```
para_I SVSESLSASQWTSYSESLASTSVSKSDSYSQSASLLSSSESASTSMPVSEFPLTSVSDSI       1557
para_F SVSESLSASQWASHSESLASASLSKSDSYSQST--FSASESTSTSMPVSEFPLTSVSESM       1423
       ***********:*:******:*:*********:  :*:***:***************:*:

para_I SASSTESQSLSQEISEWISVSYASSFSAVTSSSESVVSSFGTSYSESPSDASSLTATHSS       1617
para_F SASSTESQSLSHEASEWISVSYASSFSAMTSPSESVVSSSRTSHSESPSDASSLSATHSS       1483
       ***********:* **************:** *******  **:**********:*****

para_I GPALPETGAHPSSNILATGASILLSGLALLGIRKKDGK  1655
para_F GPALPETGAHPSSNILATVVSILLSGLALLGILKKDGK  1521
       ****************** .************ *****
```

| Protein: | accessory Sec-dependent serine-rich glycoprotein adhesin |
|---|---|
| **Gene location in *S. parasanguinis I* genome:** | NODE_12_length_79046_cov_686.224962;76950;79046;- |
| **Gene location in *S. parasanguinis F* genome:** | NODE_27_length_5522_cov_308.585124;4470;5522;- |

| Color code | Protein name | Location (para_I) | Location (para_F) |
|---|---|---|---|
| Color | MucBP domain | 104-193 / 198-286 / 299-388 / 401-488 / 499-588 | 53-140 / 151-240 |
| Color | LPXTG cell wall anchor domain | 663-698 | 308-348 |

```
para_I MSASESASVSASQSASLSTSASASASVSASQSASLSTSASESASVSASQSASLSTSASAS      60
para_F ------------------------------------------------------------      0


para_I ASVSASQSASLSTSASTSLSTSVSQSASNSSSNSESETPKQGEVIITYVDTKGKVIKDPR      120
para_F ------------------------------------------------------------      0


para_I QDTPNSPYDTPYNTTEEGEKPNTIKTPDGKTYKIVPKGDYPVGKVDGDGHLESSDPIKGK      180
para_F ------------------------------------------------------------      0


para_I VDKPKSTITYVYKEVKGNVYVHYVDVNGNKIKESVTDEKDQPVDKDYDTVVDNRPSTIEF      240
para_F ------------------------------------------------------------      0


para_I QGKTYELVPAGNYPVGKVDDQGHWTGDDATTGKVAEEDTNVTYVYQLKEDPTKPKEGDVI      300
para_F ------------------------------------------------------------      0


para_I ITYVDENGKEIQKPRQDTPNSPYDTPYNTTEEGEKPNTIKTPDGKTYKIVPKGDYPVGKV      360
para_F ------------------------------------------------MVPKGDYPVGKV      12
                                                       :***********

para_I DGDGHLESSDPIKGKVDKPRSIITYVYKEVKEEPTQPKGSVYVHYKDTEGNTIKESVTDE      420
para_F DGDGHLESSDPIKGKVDKPRSIITYVYKEVKEEPTQPKGSVYVHYKDTEGNTIKQSVTDE      72
       ******************************************************:*****

para_I LDQPVGKDYNTVEDNRPQYIRFEGKTYEIVPVGNYTVGKVDTQGHLESTDPTTGKVVEGR      480
para_F LDQPVGKDYNTVEDNRPQYIKFEGKTYEIVPVGNYTVGKVDTQGHLESTDPTTGKVVEGR      132
       ********************:***************************************

para_I KDVTYIYKLVEEPVQPKGNVYVHYVDENGNTIKTSVVDEKDQPVGKDYDTVVDNRPKTIT      540
para_F KDVTYIYKLVEEPVQPKGNVYVHYVDENGNTIKTSVVDEKDQPVGKDYDTVVDNRPKTIT      192
       ************************************************************

para_I TADGKVYELVPQGNYPVGNVDGEGHLTTTDPTTGKVIEGDKNVTYVYKLVKTPNVPTPNT      600
para_F TADGKVYELVPQGNYPVGKVDGEGHLTTTDPTTGKVIEGDKNVTYVYKLVKTPNVPTPNT      252
       ******************:*****************************************

para_I PVPPTPTPNTPVPPTPTPNTPVDPTPNKPMDPTPNTPVDPTPNTPVNPVPEQPAKPAPAL      660
para_F PVPPTPTPNTPVPPTPTPNTPVDPTPNKPMDPTPNTPVDPTPNTPVNPVSEQPAQPAPAL      312
       ************************************************* ****:*****
```

```
para_I EQLPNTGETGSVASALLGAVAGVAGVAALGSRKKEDEK  698
para_F EQLPNTGETGSVASALLGAVAGVAGVAALGRRKKEDEK  350
       ****************************** *******
```

| | |
|---|---|
| **Protein:** | SspB-related isopeptide-forming adhesin / FctA domain-containing protein |
| **Gene location in *S. parasanguinis I* genome:** | NODE_14_length_68607_cov_680.648606;19625;29671;- |
| **Gene location in *S. parasanguinis F* genome:** | NODE_19_length_41685_cov_317.811551;9715;21759;- |

| **Color code** | **Protein name** | **Location (para_I)** | **Location (para_F)** |
|---|---|---|---|
| Color | YSIRK Gram-positive signal peptide | 6-30 | 6-30 |
| Color | Streptococcal pilin isopeptide linkage domain | 702-813 / 825-943 / 1164-1282 / 1504-1608 / 1831-1949 / 2170-2276 / 2498-2602 / 2825-2930 / 3149-3252 | 705-820 / 831-935 / 944-1046 / 1057-1161 / 1170-1272 / 1283-1385 / 1394-1496 / 1506-1609 / 1843-1948 / 2170-2288 / 2509-2615 / 2837-2941 / 3164-3268 / 3491-3596 / 3815-3918 |
| | Streptococcal pilin isopeptide linker superfamily | 678-815 / 816-943 / 1155-1283 / 1494-1610 / 1821-1950 / 2163-2277 / 2488-2604 / 2815-2931 / 3140-3254 | 695-822 / 823-936 / 937-1047 / 1048-1162 / 1163-1274 / 1275-1387 / 1388-1498 / 1499-1616 / 1834-1949 / 2160-2289 / 2502-2616 / 2827-2943 / 3154-3270 / 3481-3597 / 3806-3920 |
| Color | Adhesin isopeptide-forming adherence domain | 993-1137 / 1331-1483 / 1661-1810 / 2001-2150 / 2328-2477 / 2655-2804 / 2982-3127 | 1676-1822 / 2000-2149 / 2337-2488 / 2667-2816 / 2991-3143 / 3321-3470 / 3648-3793 |
| Color | LPXTG cell wall anchor domain | 3308-3348 | 3974-4014 |

```
para_I MKDVFNKRQRFSLRKYSVGVCSVLLGTALFAAGANTASAAETTASSDASTSASTESASDS      60
para_F MKDIFNRRQRFSLRKYSIGVCSVLLGTALFAAGAQSASADEATAASESAGTAASEAAQPA      60
       ***:**:**********:****************::*** *:**:*::: :*::*:*. :

para_I TVATATAASSPEASYEVP-ATNVNQVDTVAAQEVKAQSEANKAAEKAEETAQPAPKAEET      119
para_F TTESS-QAEAPAASKAYGEGGSVPKIDLSGTAAA-TSETPASAIEKAEATATPAATE-KQ      117
       *. ::  *.:* **     . .* ::*  .:  . :..   .* **** ** ** .  :

para_I VKPA---KAEEAAQPAPKAEAPKATATSEAKKADEKSQAHSAASEAAKPVAS--TSAATS      174
para_F VAPAETKKTEEASKPLNVGSLPEIVLP-TAKIAETSSKP--ASTTAATPAATTATRAAAG      174
       * **   *:***::*   .. *: .    ** *: .*:   *:: **.*.*:  * **:.
```

```
para_I ESSETASASEAA-----AVALSTTVDLGSLRSADAP--TADRTAAVGPSATLDRAATDLT      227
para_F ESSERAAAREEAVTPAATTTFSATVNPAASITGTEPAAQTDKATSTDAAAAVANAATERT      234
       **** *:* * *     :.::*:**: .:  :.  *   :*::::.. :*:: .***: *

para_I NAGALATSRSRRNRRAVTNNAVTGDHNTNPVAVSTYLKDGETVDPAITNPNGATVKSQEV      287
para_F NAGALAVSSRRRGKRAL------TDHNNEPVAVETYLKDGEKATPGMKDPNGATVSSQTV      288
       ******.*  **.:**:       ***.:****.*******.. *.:.:******.** *

para_I PAGYQAKEGDWYTYSIIDLTRFNERYNTNYYTRAYKRFDDSTETTVELIDKTTGNVVETR      347
para_F PAGYAAKEGDVYTYSIVDLTRFNERYNTNYYTRAYKGFNDSTDTTVELIDKNTGNVVETR      348
       **** ***** *****:******************* *:***:********.********

para_I TLSASSGIQKFTTTTAASNGQLTVKYDYNKGLGAGPGKTDEPFIQFGYEVGASIQALVNP      407
para_F KITASSGIQKFTTTATASRGELTWQVDYDPGTGAGPGKTDQPFIQYGYEVGASIQALVAP      408
       .::***********::**.*:** : **: * ********:****:************ *

para_I KNE---AEQKLYQDVYNARTSTDIINVVEPAYNGRTITDSNAKIPKFVEKPTYYRVVDKN      464
para_F GHQLTRDEQKLYDAVYAARTSTDIINVVEPAYNGRTITDTNAKIPASVNKTTYYKVVDKN      468
        ::    *****: ** **********************:*****  *:* ***:*****

para_I NATFNANKTDKTVQDYVPNGNEVDLAKYATKAMEGQHFTASGERQFDGYKLYQTANPDST      524
para_F NPTFNANKTDKTVQDYVANGNEVDLASYTLKAMEGQDFTASGERQFDGYKLYQAADANDQ      528
       * *************** ********.*: ******.****************:*: :.

para_I TGFVSRPYVVGTKFMDAERAGIKRIKEIVGEDGSVVVRVYLLDPKQQSKRSDGTLSTDGY      584
para_F SGYVSRPYKVGTKFMDAERAGIKRIKEIVGEDGTVVVRVYLLDPKQQSKRSDGTLSTDGY      588
       :*:***** ************************:**************************

para_I MLLAETKPIKPGEYNTQDLVVKKSPLNTIAFTDNKGVNHPNGVEVPFDFQTAAGYTPKKT      644
para_F MLLAETKPIKPGDYNKQELNVKKSPLNTIPFTDSKGVTYANGKEVPFDFQKAAGYTPYKT      648
       ************:**.*:* ********* ***.***.: ** *******.****** **

para_I VFVPFLGDGIGHLSPNSQLENGAYVQIGTNVDLLNSLTPYKPTVYYYVKQEPVEVTPEVE      704
para_F VFVPFLGDNIGHLSPNEQLVRGV-NGIGTNVDLLNSLTPYKQPIYYYVKQKPVEVTPEVE      707
       ********.*******.** .*.   ***************  :******:*********

para_I KQLEGRVLVDGEFTFKLTEES---SSPDKHEETVTNKDGKATFSKLTFNKTGVYTYTITE      761
para_F KQLEGRVLANGEFSFKIKEVQPNKSLPAYEETVTNKADGKATFSKLTFNKVGTYDYTITE      767
       ********.:***:**:.* .   * *  .* ...: *************.*.* *****

para_I QKGSDTNVDYDAMTVTMTVTVTENAQGDLQASVKYSGEGGFAASADDKIFNNYVVAPVKT      821
para_F IPGSDKNVDYDAMTVTMTVNVTENAQGDLQATVKYSAEGGFKSSADDKVFNNYVVAPVKT      827
         ***.*************.***********:****.**** :*****:***********

para_I KFDFSKKLAGRELKDGEFKFVLKDENGQEVETVANKKDGTVTF-----------------      864
para_F KFDFSKALAGRELKAGEFSFVLKDSDGKVIQTKTNTKAGVVAFDDLTFDNTQVGTHKYTV      887
       ****** ******* ***.*****.:*: ::* :*.* *.*:*

para_I ------------------------------------------------------------      864
para_F EEVIPENKETGMTYDTMKAEVTITVTKQGHVLKATNTLPADTEFNNTFTPVATQAQFKFT      947

para_I ------------------------------------------------------------      864
para_F KKLEGKELTKDAFTFELLENGNVIQTKQNAADGTIQFDAISYAAAGTHTYTVREKAGTDT      1007

para_I ------------------------------------------------------------      864
para_F NIDYDPMNAVVTVNVTKDAQTGLLNAAVTMPADTEFNNFAVAPVKTRFDFSKALAGRELK      1067

para_I ------------------------------------------------------------      864
para_F EGEFSFVLKDSNGKTLQTKTNTKQGVVAFDDLTFDNTQVGTHKYTVEEVIPENKETGMTY      1127
```

```
para_I ------------------------------------------------------------     864
para_F DPMKAEVTITVTKEGHVLKATNTLPADTEFNNTFTPVATQAQFKFTKKLEGKELTKDAFT     1187


para_I ------------------------------------------------------------     864
para_F FELLENGNVIQTKQNAADGTIQFDAISYAAAGTHTYTVREKAGTDTNIDYDPMNAVVTVN     1247


para_I ------------------------------------------------------------     864
para_F VTKDAQTGLLNAAVTMPADTEFNNFAVAPVKTRFDFSKALAGRELKEGEFTFVLKDANGK     1307


para_I ------------------------------------------------------------     864
para_F TLQTKTNTKQGVVAFDNLTFDNTQVGVHKYTVEEVQGSEAGMTYDPMKAEVTITVTKEGH     1367


para_I ------------------------------------------------------------     864
para_F VLKATNALPADTEFNNTFTPAATQAQFKFTKRLEGKELTKDAFTFELLENGNVIQTKKNA     1427


para_I ------------------------------------------------------------     864
para_F ADGSITFDAIEYNAVGEHTYTVREVAGADTNIDYDSMNAVVTVNVTKNAATGILSAAVTM     1487


para_I -----------------------------------------------------------T     865
para_F PEDTEFNNYVVSPVVTKFDFTKKLAGRKLAAGEFSFVLKDAAGNKVETVKNDADGNVTFS     1547
                                                                  :

para_I EISFDNTKVGTHTYTVEEVIPATKEVGMTYDTMKATITVEVAKNGHALTTVTNVSSTGGV     925
para_F ELSFDNTKVGTHTYTVEEVIPANKEFGMTYDQMKATVTVEVAKNGHSLTTVTNVTSTGGK     1607
       *:********************.**.***** ****:*********:*******:****

para_I DANGNATDGTADKEFNNKITPPETPEFQPEKFVLNKEKFDLTGTKLMDDDDELQDEYTET     985
para_F DANGNATDGTPDKEFNNKVTPPETPKFQPEKFVVSKEKYDITGNKLMDDDDELTNEYTET     1667
       ********** *******:******:*******:.***:*:**.********* :*****

para_I NANPYADQVKNNEAENINTKTVERGDKLVYQVWLDTKNFTDKNNIQAVGISDTYDADKLT     1045
para_F NADPYVDKTTNNEPENLNTKTVKRGSKLVYQVWLDTTKFTEANNIQYVGVSDTYDADKLD     1727
       **:**.*:..*** **:*****:**.**********.:**: **** **:*********

para_I VNTADIKAYDSVTGVDVTSKFDITVANGVITATSKSSMNKSLGDADNTQVIDTTKFAFGR     1105
para_F VNAADIKAYDSVTGAEVTNKFDIKVENGTITATSKDEF---IKDKVNAPVIDTTKFEFGR     1784
       **:***********.:**.****.* **.******..:   : *  *: ******* ***

para_I YYKFDIPATVKADVPGGVDIENKANQIVHVYNPVSKSVETPEKPTQKRVNSVPITAEFNF     1165
para_F YYKFDIPATVKESVKAGADIENTANQTVHVYNPVSKTVEKPEKPTQKRVNSVPVPVEMNF     1844
       *********** .* .*.****.*** *********:**.*************: .*:**

para_I TKRLEGRALTAGEFTFELKDSDNVVIATATNDADGKIKFSPVEYTNKAGEKVTALKYKKG     1225
para_F TKRLEGRELQKNEFEFVLKKD-GVEVERVKNDAAGKIVFKTLEFGRD------------D     1891
       ******* *  .** * **.. .* :  ..*** *** *. :*: ..            .

para_I QEGIYTYSVTEVKGTDATVTYDTMKAEVTVTVSHDGTAKALIANVTEPADKEFNNTVTPP     1285
para_F LGKTYNYTVEETPGTDATVKYDTMVATVKVVVSHDGTAKAIVANVTDAADKEFNNRVTPP     1951
           *.*:* *. ******.**** * * *.*.*********::****: ******* ****

para_I TEPKFQPEKYVLNTAKYSITDNKLLDDDAELTDKYGETNTDPYVDKTNNNEAENINTKTV     1345
para_F EEPKFQPEKYVVSKEKYDITGDKLVDDDRELADKYADTNANPYADDASNNEAENLNTKTV     2011
        **********:.. **.**.:**:*** **:***.:**::**.*.:.******:*****

para_I NRGDKLYYQVWLDTTKFSATNKENVQSVGITDDFDETKVDVDGSAIKAYDSVTGDDVTNK     1405
para_F ERGSKLVYQVWLDTTKFDAANKDNIQTVGISDNYDEAKLNLNKADIKAYDSVTGAEVTDK     2071
       :**.** **********.*:**:*:*:***:*::**:*:::: : ********* :**:*
```

```
para_I FDIKVENGVMTATLKAGFTKSLGDAENTQIIDTTKFAFGRYYKFDIPATVKADVPGGSDI      1465
para_F FDIAVNNGVITANLKAGFTKSLGDAENTQVIDTTKFAFGRYYKFDIPTTVKDDVVAGADI      2131
       *** *:***:**.****************:*****************:*** ** .*:**

para_I ENTAAQVVNYYNPVSKTVEKPSKPTEKRVNNVPVEVEFNFTKRLEGRELKANEFSFVLKD      1525
para_F ENTAAQVVNYYNPTTKKVEKPEKPTEKRVNNVPISVEFNFTKKLEGRELKANEFTFELKD      2191
       *************.:*.****.***********:.*******:***********:* ***

para_I STGKVVETVSNDKDGNVKFSA-------------LTFKKGEEGVHNYTVEEVAGTDATVT      1572
para_F SDNVVIATATNDADGNFKFTPVDYTNKAGKTVTALKYQKGQEGTYTYTVTEVKGTDSTVA      2251
       * . *: *.:** ***.**:              *.::**:**.:.*** ** ***:**:

para_I YDTMKATVAITVEHKGTAKVLVAKLGEIADKEFNNRVTPPTEPKFQPEKYVVSKEKYDIT      1632
para_F YDPMAAVVTVKVSHDGTAKALITNVTEPADKEFNNRVTPPTEPKFQPEKYVVSKAKFDIT      2311
  ** * *.*::.*.*.****.*:::: * ************************** *:***

para_I GDKLVDDDRELADKYADTNANPYADDASNNEAENLNTKTVERGSKLVYQVWLDTTKFDTA      1692
para_F GTKLVDDDSELTDKYGETNTNPYVDNTNNNEAENLNTKTVERGSKLYYQVWLDTTKFDAA      2371
       * ****** **:***.:**:***.*::.****************** ***********:*

para_I NKDNIQTVGISDNYDEAKLNLNTADIKAYDSVTGAEVTDKFDITVNNGVITANLKAGFTK      1752
para_F NKDNIQTVGITDNYDKDKLTVNASDIKVYDSVTGADVTTKFDISDNNGVLTANLKAGFTK      2431
       **********:****: **.:*::***.*******:** ****: ****:**********

para_I SLGDAENTQVIDTTKFAFGRYYKFDIPTTVKDDVVAGADIENTAAQVVNYYNPTTKKVEK      1812
para_F SLGDTENTQIIDTTKFEFGRYYKFDIPATVKDDVVAGADIENKAAQVVNYYNPVSKKVEK      2491
       ****:****:****** **********:**************.**********.:*****

para_I PNKPTEKRVNNVPISVEFNFTKKLEGRALKANEFTFELKDSDNVVIATATNDANGNFKFT      1872
para_F PNKPTEKRVNSVPVPLDLKFTKSLEGRQLKDQEFTFVLKKDGNVV-ETVKNDATGKVNFT      2550
       **********.**: ::::***.**** ** :**** **...***  *..***.*:.:**

para_I PVDYTNKAGETVTALKYKKGQEGTYKYTVTEVKGTDATVEYDKMAAVVTVTVSHDGTAKA      1932
para_F QLKFGK------------DDLGKTYNYTVEEVRGTDSTVSYDPMVATVKVVVSHDGTAKA      2598
        :.: :            ..   **:*** **:***:**.** *.*.*.*.*********

para_I LITNVTEPADKEFNNTVRPPEEPKFQPEKYVVSKAKFDITGTKLVDDDSELTDKYGETNT      1992
para_F IVANVTDAADKEFNNRVTPPEEPKFQPEKYVLNAEKFSITDNKLLDDDSELADKYADTNA      2658
       :::***: ******* * *************:.  **.**..**:******:***.:**:

para_I NPYVDTTANNEDENLNTKPVERGQKLYYQVWLDTTKFSATNKENIQTVGITDNYDKDKLT      2052
para_F NPYVDGTANNEAENINTKTVKRGDKLVYQVWLDTTKFDAANKDNIQSVGISDDYDEAKLD      2718
       ***** ***** **:*** *:**:** **********.*:**:***:***:*:**: **

para_I VNASDIKVYDSVTGADVTTKFDISDNNGVLTANLKDGFTKSLGDAENTQIIDTTKFEFGR      2112
para_F LDSTKIKAYDSVTGAEVTDKFDIAVNNGVITATLKDGFTKSLGDAENTQIIDTTKFAFGR      2778
       ::::.**.*******:** ****: ****:**.*********************** ***

para_I YYKFDIPATVKDDVVAGADIENKAAQVVNYYNPVSKTVEKPNKPTEKRVNSVPVPLDLKF      2172
para_F YYKFDIPTTVKADVPGGVDIENTAAQVVNYYNPTTKKVEKPSKPTEKRVNNVPVEVEFNF      2838
       *******:*** ** .*.****.**********.:*.****.********.*** ::::*

para_I TKSLEGRQLKDQEFTFVLKK-DGNVVETVKNDATGKVNFTQLKFGKDDLGKTYNYTVEEV      2231
para_F TKRLEGRELKANEFSFVLKDSTGKEVETVSNDASGNVKFKALEFKKGDEG-VHNYTVEEV      2897
       ** ****:** :**:****.  *: ****.***:*:*:*. *:* *.* * .:*******

para_I RGTDSTVSYDPMVATVKVVVSHDGTAKAIVANVTDAADKEFNNRVTPPEEPKFQPEKYVL      2291
para_F KGSDATVTYDTMKANVTVTVKHDGTAKVLVATVGDIADKEFNNRVTPPEEPKFQPEKYVV      2957
       :*:*:**:** * *.*.*.*.******.:**.* * ***********************:

para_I NAEKFSITDNKLLDDDSELADKYADTNANPYVDGTANNEAENINTKTVNRGDKLVYQVWL      2351
para_F SEEKFDITGDKLVDDDKELADKYADTNANPYADKDTNNEAENINTKTVNRGDKLVYQVWL      3017
       . ***.**.:**:***.**************.*  :************************
```

```
para_I DTTKFDAANKDNIQSVGISDDYDETKLELDSTKIKAYDSVTGAEVTDKFDIAVNNGVITA      2411
para_F DTTKFSATNKENIQSVGISDDYDEAKLELDSTKIKAYDSITGAEVTDKFDIAVNNGVITA      3077
       *****.*:**:*************:**************:********************

para_I TLKDGFTKSLGDADNTQVIDTTKFTFGRYYKFDIPTTVKADVPGGVDIENTAAQVVNYYN      2471
para_F TLKDGFTKSLGDAENTQVIDTTKFEFGRYYKFDIPTTVKADVPGGVDIENTAAQVVNYYN      3137
       *************:********** ***********************************

para_I PTTKKVEKPSKPTEKRVNNVPVEVEFNFTKRLEGRELKANEFSFVLKDSEGKTLETVSND      2531
para_F PTTKKVEKPNKPTEKRVNSVPVEVEFNFTKRMEGRELKANEFSFVLKDSEGKTLETVKND      3197
       *********.********.************:*************************.**

para_I ASGNVKFSAMSFKKGQEGVHNYTVEEVKGSDATVTYDTMKANVTVTVKHDGTAKVLVATV      2591
para_F ASGNVKFSKLEFKKGQEGVHNYTVEEVKGSDATVTYDTMKANVTVTVKHDGTAKVLVATV      3257
       ******** :.*************************************************

para_I GDIADKEFNNRVTPPETPEFNPEKYVLNEKEFDLTGTSLLDDDKELADKYADTNANPYVD      2651
para_F GDIADKEFNNRVTPPETPEFNPEKYVLNEKEFDLTGTSLLDDDKELADKYADTNANPYVD      3317
       ************************************************************

para_I DASNNEAANINTKTVKPKDKLVYQVWLDTTKFDANNKDNIQSVGITDDYDEAKVDVEASA      2711
para_F DASNNEAANINTKTVKPKDKLVYQVWLDTTKFDANNKDNIQSVGITDDYDEAKVDVEASA      3377
       ************************************************************

para_I IKAYDGKTGADVTAKFDITVNNGVITATLKDGFTKSLGDAENTQVIDTTKFEFGRYYKFD      2771
para_F IKAYDGKTGADVTAKFDITVNNGVITATLKDGFTKSLGDSENTQVIDTTKFEFGRYYKFD      3437
       ***************************************:********************

para_I IPATVKADVKGGVDIENTAAQVVNYYNPTTKKVERPEKPTEKRVNSVPVSVEFNFTKKLE      2831
para_F IPATVKADVKGGVDIENTAAQVVNYYNPTTKKVEKPNKPTEKRVNSVPVQVEFNFTKRLE      3497
       **********************************:*:************.*******:**

para_I GRELKANEFSFVLKDSTGKVVETVKNDAAGNVKFSKLEFKKGQEGVHNYTVEEVKGTDGT      2891
para_F GRKLQANEFSFVLKDSEGKVVETVSNDASGNVKFSKLEFKKEQVGEHKFTVEEVKGTDAT      3557
       **:*:*********** *******.***:************ * * *::*********.*

para_I VTYDAMKAVVTVEVKHDGTAKALITNVTDPADKEFNNTVRPPETPDFNPEKYILNESKFD      2951
para_F VTYDTMKANVTVTVSHDGKAKVLIATVGDIADKEFNNKVTPPETPEFNPEKYILNESKFD      3617
       ****:*** *** *.***.**.**:.* * *******.* *****:**************

para_I LTGVKLLDDDNELEHKVADTNANPYADGTANNEAQNINTKTLKKGDKVYYQVWLDTTKFT      3011
para_F LTGVKLLDDDNELEHKVADTNANPYVDGTANNEAQNINTKTLKKGDKVYYQVWLDTTKFT      3677
       *************************.**********************************

para_I EAHNIQSVGVTDKYDSANLNVNGADIKAYDSVTGEDVTAKFDIKVENGVITATSKADLTK      3071
para_F EAHNIQSVGVTDKYDSANLNVNGADIKAYDSVTGEDVTAKFDIKVENGVITATSKADLTK      3737
       ************************************************************

para_I SLGDAENTQVIDTTKLAFGRYYKFEIPAEIKQSAQDGVDIENTASQIVHQYDPTKKTVEK      3131
para_F SLGDAENTQVIDTTKLAFGRYYKFEIPAEIKQSAQDGVDIENTASQIVHQYDPTKKTVEK      3797
       ************************************************************

para_I PEKPTEKRVVNIPVKVQFQFTKKLEGRALKAGEFSFVLKDEKGNVIETVTNDAAGKITFS      3191
para_F PEKPTEKRVVNIPVKVQFQFTKKLEGRALKAGEFSFVLKDEKGNVIETVTNDAAGKITFS      3857
       ************************************************************

para_I NLEFKRGEEGTHLYHVEEIRGTDSSVVYDKMVATVGIVVNKAGKVLVATTKLPEDTEFNN      3251
para_F NLEFKRGEEGTHLYHVEEIRGTDSSVVYDKMVATVGIVVNKAGKVLVATTKLPEDTEFNN      3917
       ************************************************************

para_I TVIPPTPPTPPTPPTPPTPPTPPTPPTPPTPPTPPTPPTPPTPPTPPTPPTPPTPTTPPA      3311
para_F TVIPPTPPTPPTPPTPPTPPTPPTPPTPPTPPTPPTPPTPPTPPTPPTPPTPPTPTTPPA      3977
       ************************************************************
```

```
para_I PALPETGEEQSASAALLGAALGMVGLAGLAKRKKRED   3348
para_F PALPETGEEQSASAALLGAALGMVGLAGLAKRKKRED   4014
       *************************************
```

| | |
|---|---|
| **Protein:** | accessory Sec-dependent serine-rich glycoprotein adhesin |
| **Gene location in *S. parasanguinis I* genome:** | NODE_14_length_68607_cov_680.648606;3;4025;- |
| **Gene location in *S. parasanguinis F* genome:** | NODE_25_length_11490_cov_302.254710;5954;11488;+ |

| **Color code** | **Protein name** | **Location (para_I)** | **Location (para_F)** |
|---|---|---|---|
| Color | KxYKxGKxW signal peptide | 14-52 | 14-52 |
| Color | Serine-rich repeat adhesion glycoprotein, N-terminal domain | 36-85 | 36-85 |
| Color | MucBP domain | 753-841 / 851-940 / 945-1032 | 778-873 / 883-972 / 977-1064 / 1614-1703 / 1717-1806 |
| | Serine-Rich Repeat Protein (SRRP) | 120-1334 | |
| Color | Overlapping domains | | |

```
para_I MFFKRSNGEFRETDRVTRFKLIKSGKNWLRAATSNFGLLKVIRGQVEETVVAEVREDAVS      60
para_F MFFKRSNGEFRETDRVTRFKLIKSGKNWLRAATSNFGLLKVIRGQVEETVVAEVREDAVS      60
       ************************************************************

para_I VKEMTSRGLLKGIVAAGAVFGAATVANTAKADETGSDVATASELSSEALVEQGSTVLGTT      120
para_F VKEMTSRGLLKGIVAAGAVFGAATVANTAKADETGSDVATASELSSEALVVQGSTVLGTT      120
       ************************************************** *********

para_I STTESQSESTTESTTESTT----ESTTESASASASASASTSTSVSVSHSASLSEQGSAEL      176
para_F STTESQSESTTASTTESTTESQSQSTTESASASASASASTSTSVSVSHSASLSEQGSAEL      180
       *********** *******    :************************************

para_I SAASSESTVAAGSEASVESTTTVAKAEDKVVLEQNTSEAALLNKIAGDYSATVSAPEKKA      236
para_F SAASSESTVAAGSEASVESTTTVAKAEDKVVLEQNTSEAALLNKIAGDYSATVSAPEKKA      240
       ************************************************************

para_I ALDAAIAKVQTELTASNSLINANASAQSYADQRERLSKSVDDMMATLTAAGFTGNTTVNG      296
para_F ALDAAIAKVQTELTASNSLINANASAQSYADQRERLSKSVDDMMATLTAAGFTGNTTVNG      300
       ************************************************************

para_I APAISAQLAPIATSNTLAAGVDATPGMDDANGATLTDKATSIPSGYAADPAANRMTFGVW      356
para_F APAISAQLAPISTSTT--GSVDTTPVITNANGATIEDAA-FNKSGYALDPNADRFTFGVW      357
       ***********:**.*  ..**:** : :*****: * *    **** ** *:*:*****

para_I NLKSYNQ----DYNTNYYVTLSVDKTT----STNNPDVYVRLVDKNTGSEVASTTLSSAN      408
para_F QFLKTNHTTGAKTNFDYYATLSVDRSAITGSLSANPDVYLRIVKKSDGSETYTRTIHAGE      417
       :: . *:    . * :**.*****:::     : *****:*:*.*. ***. : *: :.:

para_I AGKEVNLGNLSSK---TYYPYLVYNAS--TDSGSASVDIIIKNNGNVSFAKTTEQVYDYL      463
para_F SNI--SLPNYITNNNSDVTNTLTYTGANGTDPGNVTFSLV---NDN--LEYETLQIRDTN      470
       :.   .* *  ::        *.*..:  ** *..:..::   *.*  :   * *: *

para_I TPANQTESKPNVEIGVPSTKMSQTTHYKVVDTASSTYNANRSTADTTTQSYKPTGNETEL      523
para_F YPGNENEDKPNVQTSVPYAKADQTTYYKVVDSSK----------YTKGQTYTPTGNETVL      520
        *.*:.*.****: .** :* .***:*****::.           *. *:*.****** *
```

```
para_I ASYTQTGIQGQNYTASNPRSFEGYVLYQQADA--STMSGELGN-SVGTKYAELKGTRQHY      580
para_F ASYTQTGLAGQQFTASGNRNIEGYEQVPATTDTTQKTSGILGKGVVGQKLVELQGGANHY      580
       *******: **::***. *.:***     :    .. ** **:  ** * .**:*  :**

para_I YVKRIREVVANDGSTVTKLYVLDPSSVSTYNEATMGNNTDTTGYTLVYTTPVIKPGEKYI      640
para_F YVKRISEVVDTNGSTVTKLYALDPSQVSNFSASDVGT-EDVSKYTLIYTSSVNKAGDTWN      639
       ***** *** .:********.****.**.:. : :*.  *.: ***:**: * * *:.:

para_I PSATSMDTDKVLVSSKNGDYQIQVSPWHNPDKPEEGLVYLSGWYTAGHHGEKAYMFLEHP      700
para_F STGS----KTTRVNSKNGDYYIDVVETAG-----SNSLVITGWQSTTE---KVYLTHDET      687
        :.:    ... *.****** *:*    .     .. : ::** :: .   *.*:  :.

para_I KGT---------------ESSTPGGNKENVVGGQNS-----------------------G      722
para_F KVTEGFDKPFTGAPDVSLTPAGKGNNKWNLIAGRNADIVAKDKVTDSTGKVSYKELKGFT      747
       * *                 :  *.** *::.*:*:

para_I NFGNQFSILSANEKPSGDTVHYYKKTDEKGNVYVHYKDTEGTTIKASVTDEDKQPINKAY      782
para_F SFSNNYSV-PSAVKPSTDVNYYFVKSDEKGSVYVHYRDTEGNEIKASVTDEDKQPINKKY      806
       .*.*::*:  :  *** *.  :*: *:****.*****:****. **************  *

para_I DTVVDNRPATIEYNGKTYELVPAGTYTVGQVDSDGHLTT--------SDDVKGSVAKEDK      834
para_F DTVVDNRPDTIEYNGKTYERVQAGDYTVGKVGDESNLVKSDLSTVKGTDNVIGTVAKQDK      866
       ******** ********** * ** ****:*..:.:*..        :*:* *:***:**

para_I NVTYIYKVKETPKEGEVVITYVDTKGNEIQKSRQDTPKSPYDTPYDTTEKGEKPNIIKTT      894
para_F NVTYIYKVKETPKEGEVVITYVDTKGNEIQKSRQDTPKSPYDTPYDTTEKGEKPNIIKTT      926
       ************************************************************

para_I DGKTYKIVPKGDYPVGDVDENGHLKSSDPITGKVDKPKSTITYVYEEVGSVFVHYKDING      954
para_F DGKTYKIVPKGDYPVGGVDENGHLKSSDPITGKVDKPKSTITYVYEEVGSVFVHYKDING      986
       ****************.*******************************************

para_I NTIMGSVIDEQDQPLEKDYDTVVDNRPKEIKFEGKTYELVEAGNYPVGQVDSQGHWTGDD      1014
para_F NTIMGSVIDEQDQPLEKDYDTVVDNRPKEIKFEGKTYELVEAGNYPVGQVDSQGHWTGDD      1046
       ************************************************************

para_I DTTGKVASGEKNVTYIYKLKVESQSDSGSNSASDTTSASQSESEVVSNSRSVSEQASTSL      1074
para_F DTTGKVASGEKNVTYIYKLKDESQSDSGSNSASDTTSESLSQ--------SKSEQASKSA      1098
       ******************** **************** * *:        * *****.*

para_I SESASTSLSQSAVESASVSASQSSSLSTSAQESASVSASQSASLSTSASASASVSASQSA      1134
para_F SESASASLSQSTVESASVSASQSASLSTSAQESASVSASQSASLSTSAQESASVSASQSA      1158
       *****:*****:***********:************************. **********

para_I SLSASASASASVSASQSASLSTSASASASVSASQSASLSTSASESASVSASQSASLSTSA      1194
para_F SLSTSASTSASTSASQSASLSTSAQESASVSASQSASLSTSASESASVSASQSASLSTSA      1218
       ***:***:***.************. **********************************

para_I SESASVSASQSASLSESASESASVSASQSASLSESASASESASQSISESQSASTSQSVSE      1254
para_F SESASVSASQSASLSTSASESASVSASQSASLSTSASASESASQSVSESQSASTSQSVSE      1278
       *************** ***************** ***********:**************

para_I STSASESASVSASTSTSESVSASESASASLSASASESASVSASQSASLSTSASASASTSA      1314
para_F STSASESVSESASTSTSESVSASESASASLSASASESASVSASQSASLSASASESASVSA      1338
       *******.* ***************************************:*** ***.**

para_I SQSASLSASASESASVSASQSASLSTS---------------------------------      1341
para_F SQSASLSTSASASASVSASQSASLSTSASASASVSASQSASLSTSASESASVSASQSASL      1398
       *******:*** ***************

para_I ------------------------------------------------------------      1341
para_F STSASESASVSASQSASLSTSASASASTSASQSASLSTSASASESASQSVSESQSASTSQ      1458
```

```
para_I ------------------------------------------------------------      1341
para_F SVSESTSASESASESASTSTSESVSASESASASLSASASESASVSASQSASLSTSASESA      1518

para_I ------------------------------------------------------------      1341
para_F SVSASQSASLSTSASASASTSASQSASLSTSASESASVSASQSASLSTSASESASVSASQ      1578

para_I ------------------------------------------------------------      1341
para_F SASLSTSASTSLSTSVSQSASNSSSNSESEKPQGEVIITYIRENDGKEIKVQRQDTPKSD      1638

para_I ------------------------------------------------------------      1341
para_F YNTPYDTTENDEQPKYIEFEGKKYERVPAGDYPVGKVDSEGHLETSDPIKGKVEKPVSKI      1698

para_I ------------------------------------------------------------      1341
para_F TYVYKEVKEDPTKPKEGDVIITYVDENGKEIQKPRQDTPNSPYDTPYNTTEEGEKPNTIK      1758

para_I ------------------------------------------------------------      1341
para_F TPDGKTYKIVPKGDYPVGKVDGDGHLESSDPIKGKVDKPRSIITYVYKEVKEDPTKPKEG      1818

para_I ---------------------------      1341
para_F DVIITYVDENGKEIQKPRQDTPNSPYD      1845
```

| | |
|---|---|
| **Protein:** | GBS Bsp-like repeat-containing protein |
| **Gene location in *S. parasanguinis I* genome:** | NODE_16_length_62024_cov_710.654366;32435;35197;+ |
| **Gene location in *S. parasanguinis F* genome:** | NODE_9_length_95260_cov_325.969543;36405;38858;+ |

| Color code | Protein name | Location (para_I) | Location (para_F) |
|---|---|---|---|
| Color | GBS Bsp-like | 136-223 / 239-326 / 342-423 / 445-533 / 548-637 / 653-739 | 141-224 / 242-326 / 344-428 / 445-534 / 550-636 |
| Color | CHAP domain | 808-894 | 705-791 |
| Color | Papain-like cysteine peptidase superfamily | 845-903 | 742-800 |
| Color | Overlapping domains | | |

```
para_I MKKKDLIFYASATVLLAFSTQQVKADEQTSSDQTFEKTATIVLKAETSSNTENTGIHAER      60
para_F MKKKDLIFYAGTAVLLAFSTQQVKADEQHASDQTPENTSAIVATTTSSEAQNTV----SE      56
       **********.::*************** :**** *:*::** .: :*.  :..    ..

para_I SVAIEKKADTEAYRNETAKNNAEFAEYVAEEKIETESPSSAVFTSLSSNRKEEHTSATTS      120
para_F EVGVEKGIQTEN-------------------TVATEA--APVTPTLSN------------      83
       .*.:**  :**                    .: **:  : *  :**.

para_I GTIPTTEAKASGTLLIENNNPVAGTFDAVVRDIEAPNGLKEVLVPTWSLENGQDDLIWHK      180
para_F -----------------NQSSDKGTSDKVVSEQPAPAVV---------------------      105
                        *:.   ** * ** :  **  :

para_I AMREPDGSYRAKIKASDHKDSTGNYRADAYVIDKKGRAQYLSQKIVAVDYARPSGALSIE      240
para_F ASRAPYG-------------------ARARVVAARNES--VPQ------QESPVSADLTI      138
       * * * *                   * * *:  :..:  : *         * .*

para_I NNNTVAGTFDAVIRNIVAPNGVKEVLVPSWSLENGQEDLIWHKATKQSDGSYRVTIKATE      300
para_F AKSESDGTFTITAKNLQGLDGYEEVKIPFWSHANGMKDIIWYTPSRQADGSYTVTAKASD      198
        :.   ***  . :*: . :* :** :*  **  ** :*:**:. ::*:**** ** **::

para_I HKGNKGKYRADAYVVDNSNNRHYIAEKVVAVDYTRPRGVLSIENNDTVAGTFDAVVRDIV      360
para_F HENADGKYEAQVFYVDAKGQNKFVKKAFIDYTAPKPSADLTITK-SESDGTFTITAKNLQ      257
       *:. .***.*:.: ** ..:.::: : .:     :* . *:* : .   ***  ..:::

para_I APNGVKDILVPSWSLAGGQDDLIWHKATRQADGSYRVTIKATDHKNSTGRYRADAYLVDN      420
para_F GFDSYKEVKIPFWSHANGIKDIIWYTPTRQADGSYTVTAKASDHENADGKYEAQVFYVDA      317
       . :. *:: :* ** *.* .*:**:. ******** ** **:**:*: *:*.*:.: **

para_I SNTPFYLTEKVVEVTQTRPTASLIIENNNAELGTFDAVVRNISAPNGIKEVLVPSWSLVN      480
para_F NGQNKFVKKAFIDYTAPKPSADLTITKS-ESDGTFTITAKNLQGFDGYKEVKIPFWSHAN      376
       ..   ::.: .:: *  :*:*.* * :.  . ***  ..:*:.. :* *** :* ** .*

para_I GQDDLIWHKATRQPDGSYRVTIKSDEHKNSLGNYRADLYIVDNKNQHHYITETVVDVKH-      539
para_F GMKDIIWYTPTRQADGSYTVTAKASDHENADGQYEAQVFYVDAKGQNKFVKKAFIDFKNQ      436
       * .*:**:. *** **** ** *:.:*:*: *:*.*::: ** *.*::::.::.:*.*:

para_I NKPIGTISIVNNNKDTGTFDVIIKDVYSPKGVRTVQVPTWSDKDGQDDLRWYEATRQANG      599
para_F SRPTASLLIQNNNKDAGTFDVIIKDVYSPKGVRTVQVPTWSDKDGQDDIRWYEATRQSNG      496
       .:* .:: * *****:********************************:********:**
```

```
para_I DYKVSVKVSDHKNSTGKYFVHLYYIQNDGTRVGVGGTTTDVEFRNAKTKTQAYIKNVNSG        659
para_F DYKVSVKASDHKNSTGKYHIHLYYIQNDGSRVGVGSTTTEVEFRNAQTKTQTGIKNVNSG        556
       *******.**********.:*********:*****.***:******:****: *******

para_I AGTYTVTVDQAPQGRRIKNIRVAAWSQAHQENLFWYSTAPSGMHTEVQVSAANHQYQSGN        719
para_F AGTYTVTVDQAPQGRRIKNIRVAAWSQAHQENLFWYSTAPSGMHTEVQVSAANHQYQSGN        616
       ************************************************************

para_I YTTHVYVDYVDGGVEGFNLGQTALHPRATVDQTAFSPRVTNGQRDRVLRAAASLVGVRGG        779
para_F YTTHVYVDYVDGGVEGFNLGQTALHPRATVDQTAFSPRVTNGQRDRVLRAAASLVGVRGG        676
       ************************************************************

para_I TAAHQQLINDYNSVKPLPVGYAVKTTDDWCDIFVTTVFQREGLSGLIGRECGVERHIQIF        839
para_F TAAHQQLVNDYNSVKPLPVGYAVKTTDDWCDIFVTTVFQREGLSGLIGRECGVERHIQIF        736
       *******:****************************************************

para_I KRLGIWNEDGTTTPKAGDIITFNWDQNSQQNNGFADHIGIVESVSNGIIHTIEGNSNNQV        899
para_F KRLGIWNEDGTTTPKAGDIITFNWDQNSQQNNGFADHIGIVESVSNGIIHTIEGNSNNQV        796
       ************************************************************

para_I RRNTYRIGHGNIRGFATPRYQ     920
para_F RRNTYRIGHGNIRGFATPRYQ     817
       *********************
```

| | |
|---|---|
| **Protein:** | FctA domain-containing protein / Cna B-type domain-containing protein |
| **Gene location in *S. parasanguinis I* genome:** | NODE_3_length_133837_cov_553.245731;85084;89274;+ |
| **Gene location in *S. parasanguinis F* genome:** | NODE_14_length_66095_cov_250.109576;38452;40638;- |

| Color code | Protein name | Location (para_I) | Location (para_F) |
|---|---|---|---|
| Color | Fibrogen-binding domain 1 | 26-155 | 22-155 |
| Color | Adhesion domain superfamily | 41-152 | 34-152 |
| Color | SDR-like Ig domain | 45-130 | |
| Color | Collagen-binding surface protein Cna-like, B-type domain | 352-441 | 352-441 |
| Color | Streptococcal pilin isopeptide linkage domain | 473-568 / 582-687 / 700-809 / 827-941 / 957-1071 / 1084-1189 | |
| | Streptococcal pilin isopeptide linker superfamily | 454-571 / 572-690 / 691-811 / 814-940 / 949-1065 / 1066-1190 | |
| Color | Immunoglobulin-like fold | 1201-1306 | |
| Color | Prealbumin-like fold domain | 1207-1291 | |
| | LPXTG cell wall anchor domain | 1355-1394 | |

```
para_I MKSLYKKIVAFVAIIAVVALGLSVIKPVSAASVSPTVTNLKAQASGQKVTFSFDWDLTGK      60
para_F MKSLYKKIVAFVAIIGVVALGLSVIKPVSAATVSPTVTNLKAQASGQKVIFSFDWDLTGK      60
       ***************.***************:***************** **********

para_I SVKEGDTFTIDAPEGVNITEVATQSLQANGAEVATISMTNKKITFTFKKAIESMNENVKG      120
para_F SVKDGDTFTIDAPEGVNITEIATQSLQANGAEVATISMTNKKITFTFKKAIESMNENVKG      120
       ***:****************:***************************************

para_I GFSYNAVWDNTPGNPGNKTATSKVGSESVIITRPDGPGVFESVLNKYNLDGSYVTKQFKL      180
para_F GFSYKAEWDSTPGNPGNKTATSKVGSESVIITRPDGPGVFESVLNKYNLTGDYVAKQFKL      180
       ****:* **.*************************************** *.**:*****

para_I DASENYAWLNVGDDYYLTKWFIRINGDGKKQAITNPVVSDKIQAPAVDYSKITFAPAANH      240
para_F DASENYAWMNVGDDYYLTKWFIRINGDGKKQAITNPVVSDKIQAPAVDYSKITFAPAANH      240
       ********:***************************************************

para_I AANEFFVGTYLKPSFTLRKGGQVVASGWDFWKHVKFDADGNGFTVNLSDVSDVFKTASSD      300
para_F AANEFFVGTYLKPSFTLRKGGQVVASGWDFWKHVKFDADGNGFTVNLSDVSDVFKTASSD      300
       ************************************************************

para_I ELIVEYQTLIPKTTIRVDNNATLTADEITTPQTDPAFWNNPELKFWVSGDKTVTVQKEWV      360
para_F ELIVEYQTLIPKTTIRVDNNATLTADEITTPQTDPAFWNNTELKFWVSGDKTVTVQKEWV      360
       **************************************** *******************

para_I GDEEADRKDITVQLYADGKALDGMTQTLTKASGWTAEFSKLPGIKDGQPIVYSVEETNTP      420
para_F GDEEADRKDITVQLYADGKALDGLTQTLTKASGWKAAFTNLPGIKDGKKIEYSVVETNTP      420
       ***********************:**********.* *::*******: * *** *****
```

```
para_I DGYTSKVEPINESNVIKVVNTSNKPKVTETTANLVVKKAFEVAGDQEHTKLPITEGQFEF     480
para_F EGYTSKVEKIDDDNVIKVVNTSNKPTTTTTTTTTTTTTQEPTTTTT--TTTQEPTTTTT---     475
       :******* *::.************..* **:. ...:   .:     *: * *

para_I VLKDENKKVVETAKNQADGTVNFKSLTFNKEGTHTYTITENKGTDA-NVNYSTQSITATV     539
para_F -TTQEPTTTTTT--TQEPTTT----TTTTQEPTTTTTTTQEPTTTTTTTQEPTTTTTTTQ     528
        .:* .... *  .*   *.     * .:* * * * *::  * : ..:  * : *:*

para_I DVKKTDDKLVASVTYSGGDGEQKNTITNTQNKPKVSNAKVTLNLKKAFEGGELKGDDFEF     599
para_F EPTTTTTTTQEPTTTTTTTQES-TTTTTTTQEPTTTTTTT-------QEPSTT-----TT     575
       : ..*  .    .* :    *. .* *.* ::*..:.:..        * .

para_I VAKDANDQVVGTAKNQKNGSITFDNITVDKAGTFKYTITETKGTDKTITYSDKTITATVV     659
para_F TTQEPT-TTVTTTDEPKTTSTTTD-----EP---KTTVTTTDEPSTTSTTSEEPKTTVPT     626
       .::: .  .* *:.: *. * * *     :    * *:* *.  ..* * *::  *:. .

para_I VVEKDNALV---VEQISYSDGQTDTDTFTNKKEAPKTESVTATLQVNKLLKEGETNLPLT     716
para_F TPETPDTTPEEPGNHNSSEEGTTSTTTT--TTAEPKTT----PEKPNKPDHSGTTTTPSA     680
       . *. ::      :: * .:* *.* *   ..  ***       : **  :.* *. * :

para_I DDQFEFVLKEGNNTLETAKNKANGTVTFKELSYTAEGTHTYTITENKGTDASINYSTQTI     776
para_F PGS-----NGGN--------NGGG---------------RKTLLPNTGEV----------     702
        ..     : **        :..*                 *:  *.*

para_I TATVEVKKVNDKLVATVTYSGGDAEKGDTFTNTKTPPTPVPPTVKPTTAQFKAKKVLAIN     836
para_F --------------------------------------------VASGLVFS--GILVLA     716
                                              :   *.   :*.:

para_I GSS--DRTLKANEFTFLLKDQAGTLVDTKTNGENGDILFNPVSFNEAGTFTYTITEQKPA     894
para_F GAVGIKRKLTDN------------------------------------------------     728
       *:   .*.*. *

para_I TPESAITYDESVHTVTVTVTKDANGQLNADVQYDGKKNTPTFTNTYTPPTPVPPTVKPTS     954
para_F ------------------------------------------------------------     728

para_I AQFKAKKVLAINGTSDRTLKANEFTFLLKDQAGTLVDTKTNGENGDILFNPVSFNEAGTF     1014
para_F ------------------------------------------------------------     728

para_I TYTIVEQKPATPESAITYDETVHTVTVTVTKDENGQLNADVQYDGKKDTPTFTNTYTPPT     1074
para_F ------------------------------------------------------------     728

para_I PPTPSEKQITTSKILEGRDLKGGEFSFNLLDENGTVLQTKQNAADGTVTFDAIAYTEAMI     1134
para_F ------------------------------------------------------------     728

para_I GTYKYTIKEVVPADQANIQYDEGQVDVTVTVTKDEASNAIQAVVSYGDKKTFINKVIPPT     1194
para_F ------------------------------------------------------------     728

para_I PPTVNNPELKLYTLKVRKVDEKGDYLAGAVFGLFEADGVTPVANPYGQGQAQAISGQDGL     1254
para_F ------------------------------------------------------------     728

para_I ASFVGFEAKDYVIKELSAPSGYQLSNEVIKVSVSDYVAATNLVVDKGNVVNKLLPPPPST     1314
para_F ------------------------------------------------------------     728

para_I DIPNIPTPSNSKPKTPSPNGDKPKSNDKPKSSETPKSSDKPKENKKSLPSTGTEDHLGLL     1374
para_F ------------------------------------------------------------     728
```

```
para_I VTGLTFVATAIASMTLKKKEDF    1396
para_F ----------------------    728
```

| | |
|---|---|
| **Protein:** | CshA/CshB family fibrillar adhesin-related protein |
| **Gene location in *S. parasanguinis I* genome:** | NODE_4_length_133826_cov_513.738570;110991;119270;- |
| **Gene location in *S. parasanguinis F* genome:** | NODE_2_length_214271_cov_230.889749;90353;99331;- |

| Color code | Protein name | Location (para_I) | Location (para_F) |
|---|---|---|---|
| Color | YSIRK Gram-positive signal peptide | 9-40 | 9-40 |
| Color | Surface adhesin CshA, non-repetitive domain 2 | 222-526 | 223-524 |
| Color | GEVED domain | 629-706 | 628-704 |
| Color | CshA domain | 713-818 / 854-976 / 980-1091 / 1094-1210 / 1213-1318 / 1321-1426 / 1429-1527 / 1530-1628 / 1631-1739 / 1742-1840 / 1843-1941 / 1944-2042 / 2045-2153 / 2156-2254 / 2257-2355 / 2358-2456 / 2459-2557 | 711-817 / 853-975 / 979-1090 / 1094-1209 / 1212-1317 / 1320-1418 / 1421-1533 / 1537-1652 / 1654-1760 / 1763-1861 / 1864-1972 / 1975-2073 / 2076-2174 / 2177-2275 / 2278-2386 / 2389-2487 / 2490-2588 / 2591-2689 / 2692-2790 |
| Color | Surface protein repeat SSSPR-51 | 2560-2608 / 2609-2658 | 2794-2844 / 2844-2894 |
| Color | LPXTG cell wall anchor domain | 2718-2759 | 2951-2991 |

```
para_I MGKDLFNDRISRFSIRKLNVGVCSVLLGTLVMVGTAASAAAEEKKDTTNESVAAVATASE      60
para_F MGKDLFNDRISRFSIRKLNVGVCSVLLGTLVMVGTAASAAAEENTDTTSESVAAVATASE      60
       *******************************************:.***.***********

para_I APATSTATATSATSTAATTSTYDANAAITAPETSTAAVTSTAPASTSEASSTSTAASATS      120
para_F EPATSTATATSATSTAATTSTYDANAAITAPETSTAAATSTAPASTSEASSTSTAASATS      120
       ************************************.**********************

para_I TAAATS---ETPSLEATTTVNKAGEAASTTADKKEELASGVQAPATETPAVTAETSGGKR      177
para_F TAAATSTAAATPSLEAVTPENKATEVTSTSTDKGTQLVAGVPASS-ETSAVT-PETSGKR      178
       ******    ******.*  *** *.:**::**  :*.:** * : ** ***   :.***

para_I RNRRALGDANDPNLIGDDVEDATSTPKVEKPGFTTDLDAKSMTSQITWLDFGDVANWTGA      237
para_F RSRRSVGDPNDPNLIGDDVQDATSTPKEAKPGFTTNVKASDLASQISWLDFGDTANWTGT      238
       *.**::** **********:*******  ******::.*..::***:******.*****:

para_I KTVIVPKNDAFPNKDLEEKLALQVGATYTKEIMPGYVVTVKVKSLKPFQATEIYKKRMEE      297
para_F TTA------------SKGELALQVGATYTKEIMPGYVVTIKVKSLKPFQATEIYKKRLED      286
       .*.             : :********************:*****************:*:

para_I QGATEAEKATYDPNAKNGYVKGVTS--TGAKKAFNDGEEADVVADAQNNWTEVRHENVDT      355
para_F RGATAAEKATYDPNARNGYVNNVGGNATQARAAFNAGEEAKVIAEPQSQWTEIRKEGINT      346
       :*** **********:****:.* .  * *: *** ****.*:*: *.:***:*:*.::*
```

```
para_I KAK-KTTMGSAMNGGNIGVQFEISATFRGKTVKPAIVMADGESANPGEFVMFTTNGGGWQ     414
para_F GDKKKTTISAEFDGGNIGVQFEVSATFRGKVVKPAIVMADGESANPGELVMFTTNGEGWQ     406
         * ***:.: ::*********:*******.*****************:******* ***

para_I HLGEWKKNTRPATSVPYQPQDTDNLLGPKPKY-------NNINLRQLRDSTQVGPEKKPV     467
para_F QIGEWYKNGKTW-TRTFIPQDTDNLFGPKPTTNINGINFYTVNLTQLRGSTQAGPDKKAV     465
       ::*** ** :   :  : *******:****.         .:** ***.***.**:** *

para_I AWKYFGNPDQVTGGLGTGVFGPNISEGNYTVPLVMTRGASEVGLYIASAGKQSAMLGFFP     527
para_F AWKYFGSADLTTGGLGTGVFGPNISASDVAVPLVMTKGASEIGLYIASGGKQSAMLGFFP     525
       ******. * .************** .: :******:****:******.***********

para_I LDEGDAPESYGKAVHTIATVDGVTGAKVNQPYLGNVSPDMDENTTLDWFGDDKATTADEG     587
para_F LDEGDAPESYGKAIHSIATVDGITGKKVSQPYLGHLSPDMDENNTLDWTGDDKATTADEG     585
       *************:*:******:** **.*****::*******.**** ***********

para_I INQLLPDELKGTTNEMIKMDRTRPGNYKMSVQAHLDGASEAYIYGWVDFNQNGTFDEDER     647
para_F IDQLLPNDLKGKTNELIKMDRTRPGNYTISVEAHTGGAAKANIYGWIDFNQNGTFDEDER     645
       *:****::***.***:***********.:**:** .**::* ****:*************

para_I SELTKVTQDGTVELTFAKSKTYIDPSVNELGARVRIAKKATEIENPTGMAFSGEVEDFKT     707
para_F SDLTTITQDGTATLSFTKSKTYIDPSVKELGVRLRIAKDAAQIESPTGMAFSGEVEDFKT     705
       *:**.:*****. *:*:**********:***.*:****.*::**.***************

para_I QITHPPKGEFKETSGPQATKQTATVTFTARGEHKYEPNSHAVIDETVEPYIVDK-DGNRA     766
para_F QITHPPKGEFKETTGLQGAKQTATVAFTARGLQGYSLTEPAKIDETVAPQMIDNRTGQVV     765
       *************:* *.:******:***** : *. .. * ***** * ::*:  *: .

para_I TLDADGYYVVPGQGKYKITANGKDVDVEFIPEDNFLGTADGISIRRSDNNGYDTGWSTKF     826
para_F TPGADGYYAVAGQGKYKITPNGTSVDVEFIPEDHFLGTADGISIRRTDSNGYDTGWSTKF     825
       * .*****.* ******** **..*********:************:*.***********

para_I PDQEPNINGQLNTMDGQYVPTVTPIEIEGVDKTSTDVQGATQTGTPTFNTTATNAKGDKI     886
para_F PADEANVDTVLNTMDGLYIPTVTPTDIEGVDKTSTDVQGATQTGTPTFNTTTTNANGEKI     885
       * :* *::  ****** *:***** :*************************:***:*:**

para_I AVTPSAEYPAKLVDPATGRTTDETSVTVAGEGTYTINPSTGEVTFTPEPSFTGTAKGVDV     946
para_F SVTPSLTYPAKLVDPATGQVTNATSVTVAGEGTYSIDDATGKVTFVPEPGFTGTAQGVTV     945
       :****  ***********:.*: ***********:*: :**:***.***.*****:** *

para_I TLSAPVGRNKNGKVQEEYIKTATAKYTPTVTPITVTPTDKVSADVQNVPQTQTPTFDLSN     1006
para_F SVTAPVGRDKDGTVRDEYLKTATAKYTPTVTPITVTPTDKVSTDIQNVPQTQTPTFDLSN     1005
       :::*****:*:*.*::**:***********************:*:***************

para_I DKTAEITSKKLVDPATGQPTDETTVTVAGEGSYTIDPTTGAVTFTPEKDFVGTAKGVTVQ     1066
para_F DKTAEITSKKLVDPATGQPTDETTVTVAGEGTYTIDPTTGAVTFTPEKDFVGTATGVKVQ     1065
       *******************************:**********************.**.**

para_I ATATITNANGKTATITSDATYTPTVVPVVPTANPATSKDVQGATQTGTPTFAGTTVQVNG     1126
para_F ATATITNADGKTSTITSDASYTPTVVAAVPTANPATSKDVQGATQTGTPTFAGTTVQVNG     1125
       ********:***:******:****** .********************************

para_I EDKAITIKDNSYTLLDNDGNEVSSTPAYAEDGTTPIGTFTIDPATGQVTFTPTDKSYTGK     1186
para_F EDKAITIKDNSYTLLDKDGNEVSSTPAFAEDGTTEIGTFSIDPATGQVTFTPTDKSYTGA     1185
       ****************:**********:****** ****:******************* 

para_I VTPAKVQAESSNGIKVDTTYTPEIVPVTPTATPAETTDIQGATQTGKPEFKGGTVTVDGV     1246
para_F VTPAKVQAESSNGIKVDTTYTPEIVPVTPTATPAETTDIQGATQTGKPEFKGGTVTVDGV     1245
       ************************************************************

para_I EKTVEINEAVPAKFDDGSTTKTVEGIGTYTVAADGTVTFVPEKSFVGTAPAVTVVREDKN     1306
para_F EKTVEINEAVPATFDDRSTTKTVDGVGTYTVAADGTVTFVPEKSFVGTAPAVTVVREDKN     1305
       ************.*** ******:*:**********************************
```

```
para_I GTKASATYTPTVLPVTPTATPAETTDIQGATQKGKPEFKGGTVTVDGVEKTVEINEDVPA   1366
para_F GTKASATYTPTVTPVTPTATPAESTGVQGATQTGKPEFTAGNS-------RVPMNDDVAA   1358
       ************ **********:*.:*****.*****..*.         * :*:** *

para_I TFDDGSTTKTVEGVGTYTVAADGTVTFVPEKSFTGKAPAVTVVREDKNGTKASATYTPTV   1426
para_F TFDDGSTTKTVDGVGTYTVAADGTVTFVPDPSFTGTAPAVTVVREDKNGSKASATYTPTV   1418
       ***********:*****************: ****.*************:**********

para_I TPVT--------------------------------------------------------   1430
para_F NPVTLTPTNKVSEDIQNVPQTETPTIFALSDDETAQITSKKLIDPATGQPTDETTVTVAGE   1478
       .***

para_I ------------------------------------------------------------   1430
para_F GTYTIDPTTGAVTFTPEKDFVGTATGVKVQATATITNADGKTSTITSDASYTPTVVAAVP   1538

para_I ------------------------------------------------------------   1430
para_F TANPATSKDIQGATQTGTPTFAGTTVQVNGQDKAITIKDNSYTLLDNDGNEVTSTPAYAE   1598

para_I ----------------------------------------------------------PT   1432
para_F DGTTKIGTYSIDPATGQVTFTPTDKSYTGKVTPVKVQAESSNGIKVDTTYTPEIVPVTPT   1658
                                                                 **

para_I AKPVETTDIQGATQTGKPVFTEGD-------SRVPMNDDVPATFDNGSTTKTVDGVGTYT   1485
para_F ATPAETTDIQGATQTGKPEFKGGTVTVDGVEKTVEINEDVPATFDDGSTTKTVDGVGTYT   1718
       *.*.************** *. *         .  * :*:*******:**************

para_I VAADGTVTFVPEKSFTGTAPAVTVVREDKNGTKASATYTPTVTPVTPTATPVETTGKQGQ   1545
para_F VAADGTVTFVPEKSFVGTAPAVTVVREDVNGTKASATYTPTVTPVTPTAKDATSTGKQGQ   1778
       ***************.************ ********************.  . :******

para_I TQTGKPEFTEGDSRVPMNDDVPATFDDGSTTKSVDGVGTYTVAADGTVTFVPEKSFTGKA   1605
para_F TQTGKPEFTEGNSRVPMNDDVPATFDDGSTTKTVDGVGTYTVATDGTVTFVPEKSFTGKA   1838
       ***********:********************:**********:****************

para_I PAVTVVREDKNGTKASATYTPTVTPVTPTATPAESTGPQGLVQTGTVTFTEGDEVAPINK   1665
para_F PAVTVVREDKNGTKASATYTPTVTPVTPTATPAESTGPQGLVQTGTVTFTEGDEVAPINK   1898
       ************************************************************

para_I DSITLLDENGQPAASVEAKSPAGDVIGTYTVDKDTGVVTFTPTDKSYSGDVVPVKVQAAD   1725
para_F DSITLLDENGQPAASVEAKSPAGDVIGTYTVDKDTGVVTFTPTDKSYSGDVVPVKVQAAD   1958
       ************************************************************

para_I ANGTTVETTYTPKITPVVPTSEDATSTDIQGQTQSGKPIFTEGNPNVPIDEDTPATFEDG   1785
para_F TNGTTVETTYTPKITPVVPTSEDATSTDIQGQTQSGKPIFTEGNPNVPIDEDTPATFEDG   2018
       :***********************************************************

para_I STTKTVDGEGTYTVAPDGTVTFVPEKSFTGTATGVTVKRVDKNGTEITAKYTPTVTPVTP   1845
para_F STTKTVDGEGTYTVAPDGTVTFVPEKSFTGTASGVTVKRVDKNGTEITAKYTPTVTPVTP   2078
       ********************************:***************************

para_I TATPAESTDIQGATQTGKPKFTEGDSRVPMNDDVPATFDDGSTTKTIDGVGTYTVAADGT   1905
para_F TAEPATSTDIQGATQTGKPEFTEGDSRVPMNNDVPATFEDGSTTKTVDGVGTYTVAPDGT   2138
       ** ** *************:************:*******:*******:********* ***

para_I VTFVPEKSFVGTAPAVTVVREDKNGTKASATYTPTVTPVTPTAEDTTSTDKQGQTQTGTP   1965
para_F VTFVPEKSFVGTAPAVTVVREDMNGTKASATYTPTVTPVTPTSEDTTSTDKQGQTQTGTP   2198
       ********************** *******************:*****************

para_I TFTPGNPNVPMDDDTPATFEDGSTTKTIPGEGTYTVAPDGTVTFVPEKSFTGEGTGVTVK   2025
para_F TFTPGNPNVPMDDDTPATFEDGSTTKTIPGEGTYTVAPDGTVTFVPEKSFTGTGTGVTVK   2258
       **************************************************** *******
```

```
para_I  RVDKNGTPVTAKYTPTVTPVTPTATPAESEAPQGVVQTGTVTFTEGDPVAPIDKDTITLL      2085
para_F  RVDKNGTPVTAKYTPTVTPVTPTASPAESEAPQGVVQTGTVTFTEGDPVAPIDKDTITLL      2318
        ************************:***********************************

para_I  DENGQPAESVVAKSPEGKEIGTFTVDKETGVVTFTPKDKSYSGDVVPVKVQAKDTNGTVA      2145
para_F  DENGQPAESVVAKSPEGKEIGTFTVDKETGVVTFTPTDKSYSGDVVPVKVQGKDTNGTVA      2378
        ************************************.**************.********

para_I  ETTYTPKITPVVPTADPATSTDIQGQTQTGTPSFTPGNPAIPMDDNVPATFEDGSTTKVI      2205
para_F  ETTYTPKITPVVPTADPATSTDIQGQTQTGTPSFTPGNPAIPMDDDVPATFEDGSTTKVI      2438
        *********************************************:**************

para_I  PGEGTYTVAPNGTVTFVPEKSFTGTGTGVTVKRVDKNGTPVTATYTPTVTPVTPTAKPTT      2265
para_F  PGEGTYTVAPDGTVTFVPEKSFTGTGTGVTVKRVDKNGTPVTATYTPTVTPVTPTASPAV      2498
        **********:*********************************************.*:.

para_I  STDIQGATQTGKPEFTEGDSRVPMNDDVPATFDDGSTTKTVDGVGTYTVAPDGTVTFVPE      2325
para_F  STDVQGATQTGKPVFTEGDSRVPMNDDVPATFDDGSTTKVIPGEGTYTVAPDGTVTFVPE      2558
        ***:********* *************************.: * ****************

para_I  KSFVGTAPAVTVVREDKNGTKASATYTPTVTPVTPTATPAVSTDIQGATQTGKPVFTEGD      2385
para_F  KSFTGTGTGVTVKRVDKNGTPVTAKYTPTVTPVTPTAEPATSTDIQGQTQTGKPTFTPGN      2618
        ***.**. .*** * ***** .:*.************ **.****** ******.** *:

para_I  SRVPMNDDVPATFDDGSTTKTVKGVGTYTVAPDGTVTFVPEKSFTGTGTGVTVKRVDKNG      2445
para_F  PDVPMDDDTPATFEDGSTTKVIPGEGTYTVAPDGTVTFVPEKSFTGTGTGVTVKRVDKNG      2678
        ***:**.****:******.: * **********************************

para_I  TPITATYTPTVTPVTPTAEPATSIGKKGATQTGKPTFTEGDSRVPMNDGVPATFEDGSTT      2505
para_F  TPVTAKYTPTVTPVTPTAEPATTIGPKGKEQSGKPTFKEGDSRVPMNDKVPATFEDGSTT      2738
        **:**.****************:** **  *:*****.********** ***********

para_I  KTIPGVGTYTVAADGTVTFTPEPEFTGTAPAVTVVREDVNGTKASATYTPTVLPITKFVD      2565
para_F  KTIPGVGTYTVAADGTVTFTPEPEFTGTAPAVTVVREDVNGTKASATYTPTVLPITKFVD      2798
        ************************************************************

para_I  KDGKEIPGYPTVDGEEPKAEIPGYRFVETKKLPNGDTEHVYEKVTTSYVDENGDPIPGNP      2625
para_F  KEGKEIPGYPTVDGEEPKVEIPGYRFVETKKLPNGDTEHVYEKVTTSYVDENGDPIPGNP      2858
        *:****************.*****************************************

para_I  TEDGEQPKKDIPGYDFVKTVVDKDGNTQHIYKKTVTPTPMPDPTPTPEPQPQPTPQPQPQ      2685
para_F  TEDGEQPKKDIPGYDFVKTVVDKDGNIQHIYKKTVTPTPIPDPTPTPEPQPQPTPQPQPQ      2918
        ************************** ************:********************

para_I  PTPQPQPNPQPKPEEPTIPVVPETKEEVKYIDPQNPTAQLPNTGTKESSTAGLAIFSALA      2745
para_F  PTPQPQPTPQPKPEEPTIPVVPETKEEVKYIDPQNPTAQLPNTGTKESSTAGLAIFSALA      2978
        *******.****************************************************

para_I  GLSLFGFAKRKKED      2759
para_F  GLSLFGFAKRKKED      2992
        **************
```

| **Protein:** | GBS Bsp-like repeat-containing protein |
|---|---|
| **Gene location in *S. parasanguinis F* genome:** | NODE_9_length_95260_cov_325.969543;32793;36266;+ |
| **Gene location in *S. parasanguinis I* genome:** | NODE_16_length_62024_cov_710.654366;32435;35197;+ |

| **Color code** | **Protein name** | **Location (para_F)** | **Location (para_I)** |
|---|---|---|---|
| Color | Glycoside hydrolase, family 25 | 134-352 | |
| Color | Glycoside hydrolase superfamily | 135-341 | |
| Color | GBS Bsp-like | 364-452 / 474-557 / 579-663 / 681-765 / 782-871 / 887-973 | 136-223 / 239-326 / 342-423 / 445-533 / 548-637 / 653-739 |
| Color | CHAP domain | | 808-894 |
| Color | Papain-like cysteine peptidase superfamily | | 845-903 |
| Color | Overlapping domains | | |

```
para_F MKKKDLIFYAGAAVLMAVSAQGVSADELVSNEAATTEGNQVQAEKAPEVAVAEKSVAPVA      60
para_I MKKKDLIFYASATVLLAFSTQQVKADEQTSSDQ---------------------------      33
       **********.*:**:*.*:* *.*** .*.:

para_F SNYAAPANVTEQSVAPASKVAASESGTPSVEKATEASTTEKEETPLPSNTGSTTFFNTGA      120
para_I ------------------------------------------------------------      33

para_F HAPAGRSTDVAVQPKSFVDVSSHNGDISIGDYRTLANKGVGGVVVKLTEDTWYKNPNAEN      180
para_I ------------------------------------------------------------      33

para_F QIRNAQAAGLQVSTYHFSRYTSEEAARAEARFYIAEAQRLNIPKNTLMVNDFEDAKMQPN      240
para_I --------------------------------------------------TFEKT-----      38
                                                          **.:

para_F INRNTQAWADEMRKNGYTNLMFYTSASWLDENNLRKKGPVNTAQFGLQNFWVAQYPSPKL      300
para_I -----------------ATIV------------LKAETSSNTENTGIHAERSVAI----E      65
                        :.::            *: :   ** : *::    .

para_F SVNDAKSLRYNGKAGAWQ---FTSQA---ELLPGKHLFDHSV-DYT-GRFTANSKPAADP      352
para_I KKADTEAYRNETAKNNAEFAEYVAEEKIETESPSSAVFTSLSSNRKEEHTSATTSGTIPT      125
       .  *::: * :   .  :   :.::       *.. :*     : .  : :*.:. :

para_F TEGSLSGKIDIVNNDTMTGRFDVVISNVKAPNGVRTVSVPIWSETGGQDDLVWYTANRQA      412
para_I TEAKASGTLLIENNNPVAGTFDAVVRDIEAPNGLKEVLVPTWSLENGQDDLIWHKAMREP      185
       **.. **.: * **: ::* **.*: :::****:: * ** **  .*****:*:.* *:

para_F NGTYTVNVKAADHKNSTGLYNVHLYYVQNNGQMTGVGGTTTTVAIGKKNQTPVSADLTIA      472
para_I DGSYRAKIKASDHKDSTGNYRADAYVIDKKGRAQYLSQKIVAVDYA----RPSGALSIEN      241
       :*:* .::**:***:*** *... * ::::*:   :. . .:*  .     * .*
```

```
para_F KSEKDGTFTITAKNLQGFDGYKEVKIPFWSHANGMKDIIWYTPTRQADGSYTVTAKASDH      532
para_I NNTVAGTFDAVIRNIVAPNGVKEVLVPSWSLENGQEDLIWHKATKQSDGSYRVTIKATEH      301
       :.    ***   . :*: . :* *** :* **   ** :*:**:. *:*:**** ** **::*

para_F ENADGKYEAQVFYVDAQGQNKFVKKAFIDYTATKPANAVAADLTITKSEKDGTFTITAKN      592
para_I KGNKGKYRADAYVVDNSNNRHYIAEKVVAVDYTRPRGVLSIE---NNDTVAGTFDAVVRD      358
       :. .***.*:.: ** ..:.::: : .:     *:* ..:: :    .:.   ***   ..::

para_F LQGFDGYKEVKIPFWSHANGMKDIIWYTPTRQADGSYTVTAKASDHENADGQYEAQVFYV      652
para_I IVAPNGVKDILVPSWSLAGGQDDLIWHKATRQADGSYRVTIKATDHKNSTGRYRADAYLV      418
       : . :* *:: :* ** *.* .*:**:. ******** ** **:**:*: *:*.*:.: *

para_F DANGQNKFVKKAFIDYTASKPSADLTITKS-EKDGTFTITAKNLQGFDGYKEVKIPFWSH      711
para_I DNSNTPFYLTEKVVEVTQTRPTASLIIENNNAELGTFDAVVRNISAPNGIKEVLVPSWSL      478
       * ..   ::.: .:: * ::*:*.* * :.   : ***   ..:*:.. :* *** :* **

para_F ANGMKDIIWYTPTRQADGSYTVTAKASDHENADGQYEAQVFYVDAQGQNKFVKKAFIDYK      771
para_I VNGQDDLIWHKATRQPDGSYRVTIKSDEHKNSLGNYRADLYIVDNKNQHHYITETVVDVK      538
       .** .*:**:. *** **** ** *:.:*:*: *:*.*::: ** :.*::::.::.:* *

para_F NQSRPTGTLLIQNNNKDTGTFDVIIKDVYSPKGVQTVQVPTWSDKDGQDDIRWYEATRQA      831
para_I -HNKPIGTISIVNNNKDTGTFDVIIKDVYSPKGVRTVQVPTWSDKDGQDDLRWYEATRQA      597
        :.:* **: * *********************:***************:*********

para_F NGDYKVSVKASDHKNSTGKYHVHLYYIQNDGSRIGIGTTTTDVEVRNAMTKTQASIKNVN      891
para_I NGDYKVSVKVSDHKNSTGKYFVHLYYIQNDGTRVGVGGTTTDVEFRNAKTKTQAYIKNVN      657
       *********.**********.**********:*:*:* ******.*** ***** *****

para_F ATNGTYTVAVDQAPQGRQIKNIRVAAWSKAHQENLYWYSATPTGMHTEITVSANNHGNEA      951
para_I SGAGTYTVTVDQAPQGRRIKNIRVAAWSQAHQENLFWYSTAPSGMHTEVQVSAANHQYQS      717
       :  *****:*******:*********:******:***::*:*****: *** **   ::

para_F GNYTTHVYVDYKDGGVEGFNLGQTSLSPRNQKVNPQTTY---YSQRDP--RWAGKYYGVS      1006
para_I GNYTTHVYVDYVDGGVEGFNLGQTALHPRATVDQTAFSPRVTNGQRDRVLRAAASLVGVR      777
       *********** ************:* **     :    :     .***   * *..  **

para_F SVD-----------QSGCVPTSLAMT----FTDILGKTILPTTVAD-------------Y      1038
para_I GGTAAHQQLINDYNSVKPLPVGYAVKTTDDWCDIFVTTVFQREGLSGLIGRECGVERHIQ      837
       .                .    :*.. *:.     : **: .*::      .

para_F LYNNTDSFNKGEAGTDSDGIVAATRNWGLKSQLINGAGG---IAEALMAGK-HVLAAVGN      1094
para_I IFKRLGIWNEDGTTTPKA-GDIITFNWDQNSQQNNGFADHIGIVESVSNGIIHTIEGNSN      896
       :::. . :*:. : *  .       * **. :** ** ..    *.*::  *  *.: . .*

para_F SQFTSDPYTHELVLHGYDNGRTYVRDPYNSGNNGWYSINYLHSIKSKDPMDNKLGAPFFS      1154
para_I NQVRRNTYR-----IGHGNIRGFATPRYQ-------------------------------      920
       .*.  : *        *:.* * :.   *:

para_F IFA    1157
para_I ---    920
```
